# Pediatric traumatic brain injury elicits acute neuroinflammation and long-term changes in social, cognitive, and decision-making behaviors in male and female rats

**DOI:** 10.64898/2026.07.09.737495

**Authors:** Marissa A. Smail, Miranda Y. McDonald, Rebecca Boland, Michaela R. Breach, Courtney N. Dye, Jenna E. McCloskey, Kris M. Martens, Ashley E. Walters, Ale Zaleta Lastra, Jake Roush, Emily Yeung, Alexander Weinstein, Erin Gorman-Sandler, Cole Vonder Haar, Olga N. Kokiko-Cochran, Kathryn M. Lenz

**Affiliations:** Department of Psychology, The Ohio State University, Columbus, OH, United States; Department of Neuroscience, The Ohio State University, Columbus, OH, United States; Neuroscience Graduate Program, The Ohio State University, Columbus, OH, United States; The Institute of Brain, Behavior and Immunology, The Ohio State University, Columbus, OH, United States; The Center for Brain Injury Recovery and Discovery, The Ohio State University, Columbus, OH, United States

**Author notes:** Corresponding author: Marissa A. Smail, Ohio State University, Department of Psychology, 1835 Neil Ave, Room B85, Columbus, OH 43210, USA.

**Keywords:** pediatric, traumatic brain injury, neuroimmune, brain development, cognitive behavior

## Abstract

Traumatic brain injury (TBI) is one of the leading causes of emergency room visits in children under 10. Children are potentially more vulnerable to the adverse effects of TBI, given that their brains are still developing at the time of injury. Indeed, early life TBI has been linked to cognitive, social, and mood-related impairments later in life. The neuroimmune system has been implicated in adult TBI mechanisms and plays numerous key roles in brain development, making it an interesting candidate for linking pediatric TBI and prolonged behavioral alterations. Here we establish a rat model of mild pediatric TBI to investigate the relationship between early life TBI, acute responses of neuroimmune cells, and chronic behavioral dysregulation. At postnatal day 15, which is roughly equivalent to toddler age, male and female rat pups received a TBI via lateral fluid percussion injury. At 3 days post injury, TBI increased microglia and astrocyte coverage locally in the Perilesional Cortex but not in more distant corticolimbic regions. However, the hippocampus and prefrontal cortex did exhibit increased expression of the phagocytic marker CD68 in microglia, suggesting widespread glial activation even in the absence of gross coverage change. TBI also impacted mast cells, early-response innate immune cells, increasing their number and degranulation in multiple regions. In the juvenile and early adult periods, TBI impaired cognitive function, reduced sociability, and increased avoidance, with no change in anxiety-like behavior. Later in adulthood, TBI continued to impact cognitive behavior, increasing risky decision-making and impairing optimization months after injury. Together, these results suggest that pediatric TBI causes lasting cognitive and social dysregulation, possibly via acute neuroimmune alterations following injury at a critical period of brain development.

## INTRODUCTION

Pediatric Traumatic Brain Injury (TBI) is a major public health concern, negatively affecting over half a million children each year^1–5^. These injuries largely result from falls and impacts, along with vehicular accidents and abuse^3,5^. Pediatric TBI has an estimated lifetime incidence of 2.5%^2,6^ and annual financial burden of over 1 billion dollars^3,7^. These numbers are likely underreported, as over 80% of pediatric TBIs are mild and may not lead families to seek medical attention^5,8,9^. Beyond acute challenges, over 20% of children experience persistent behavioral symptoms after a TBI ^5,8,10,11^, including difficulties with executive function, attention, impulsivity, memory, sociability, and mood that can last through adolescence into adulthood^2,5,7,11–14^. Post-TBI symptoms can also differentially affect boys and girls^11,15–17^. Despite the established impact of pediatric TBI, little is known about the underlying mechanisms through which an injury in childhood programs persistent neurocognitive impairment throughout the lifespan^1–3^.

Pediatric TBI likely exerts lasting effects because children’s brains are rapidly developing at the time of injury^4,5,18,19^. Neuronal connections crucial to behavioral regulation are made and refined throughout childhood^20,21^, and an insult such as a TBI during this critical period could influence the formation and subsequent function of key cognitive, social, and mood-relating regions^17,21,22^. Two regions of focus in the current studies are the hippocampus (HPC) and prefrontal cortex (PFC), as both participate in diverse behavioral function^23–26^ and are particularly sensitive to early life insults^21,22^ and TBI^17,27^.

The neuroimmune system may be a key mechanism through which early life TBI shapes neurodevelopmental outcomes. Microglia and astrocytes shape development through multiple means, including synaptic pruning, cytokine release, and interactions with the neuronal environment^20,22,28,29^. Mast cells are early-response innate immune cells which communicate peripheral information centrally to trigger secondary responses, peak in number in the brain early in life, and show sensitivity to various perturbations^30–33^. As such, mast cells could also contribute to pediatric TBI outcomes. In adults, TBI causes not only acute neuroinflammation which aids in initial recovery, but also prolonged activation which can become detrimental^19,34–37^. Such heightened immune responses could have exaggerated effects during critical periods of development^21,38,39^, with aberrant activity of microglia, astrocytes, and mast cells permanently altering the structure and function of the brain.

Here, we establish a rat model of mild pediatric TBI that elicits robust acute neuroimmune and prolonged behavioral effects. We previously showed that a 1.2atm lateral fluid percussion (LFPI) in rats delivered at postnatal day (P) 15 caused mild gliosis and social dysregulation^40^, but the effects were smaller in magnitude than those seen in human behavioral studies after pediatric brain injury. Thus, in the current studies we increased the LFPI injury intensity to 2atm and examined face validity of the model via inflammatory and behavioral assessments, with the ultimate goal of utilizing this preclinical model to examine the neuroimmune system’s potential as a therapeutic candidate to ameliorate the lasting negative impact of pediatric TBI.

## MATERIALS AND METHODS

### 2.1 Subjects

Timed-pregnant Sprague Dawley rats were purchased from Inotiv. P0 was recorded as the first day pups were present. Pups were sexed between P0-3 but otherwise left undisturbed until P15 when TBI manipulations were performed. Pups were monitored daily as part of postoperative care until either euthanasia (P18) for histological assessments or weaning (P22-24) for behavioral testing. The colony room was temperature and humidity-controlled and maintained on a 12-h light-dark cycle (06:00 on, 18:00 off). All experiments complied with the National Institutes of Health Guidelines for the Care and Use of Animals and were approved by the Ohio State University Institutional Animal Care and Use Committee.

### 2.2 Experimental Design

These studies consisted of three experimental cohorts (Fig 1D). Experiment 1 was designed to assess the acute neuroimmune effects of pediatric TBI, focusing on key behavior-regulating brain regions (N = 35, 5-7 rats/sex/group). Experiment 2 examined the behavioral effects of pediatric TBI, focusing on domains known to be impacted in humans (N = 26, 4-5 rats/sex/group). Experiment 3 further interrogated long-lasting TBI cognitive outcomes utilizing more complex operant behavior (N = 31, 4-7 rats/sex/group). See Supplementary Methods for full experimental details.

**Figure 1:**
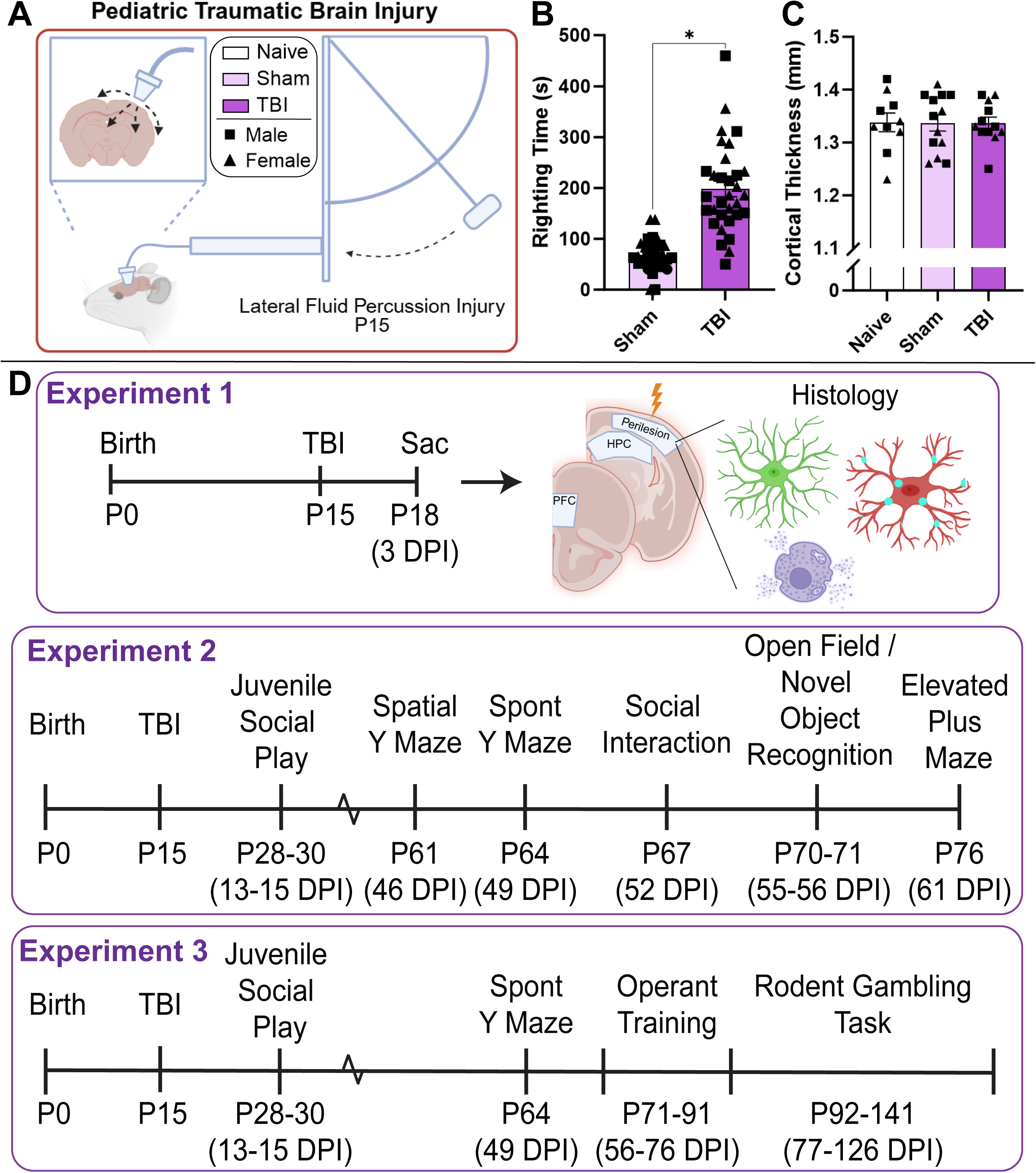
Experimental design. (A) A Lateral Fluid Percussion Injury (LFPI) device was used to deliver a mild (2atm) TBI at postnatal day (P) 15. Rats were randomly assigned to 3 groups: Naïve received no manipulations, Sham received surgery but no TBI, and TBI received a surgery and TBI. Males and females were included in all experiments. (B-C) The present model induced righting times of ∼3-5 min but did not cause cortical tissue loss. (D) Experiments consisted of 3 cohorts. The first was sacrificed at 3 days post injury (DPI) to assess acute neuroinflammatory responses. The second underwent cognitive, social, and mood-related behavioral testing in juvenile and early adult periods. The third received social play and spontaneous Y maze testing, followed by the Rodent Gambling Task in adulthood. *p<0.05. N=14-17/sex/group. Error bars= Mean±SEM. Males are denoted by squares, while females are denoted by triangles.

### 2.3 TBI Procedures

On P15, which is roughly equivalent to toddler-age in humans^20,38^, male and female pups from each litter were randomly assigned to one of three groups: Naïve, Sham, or TBI (1-3 pups/sex/litter/group). TBI consisted of a LFPI (Fig 1A; 2atm), using a protocol modified from mouse studies and optimized on P15 rats^40^. Naïve pups were left undisturbed with the dam, Sham pups received all surgical procedures except for the injury, and TBI pups received all surgical procedures and the injury. The inclusion of both Sham and Naïve control groups allows for the dissociation of effects of injury specifically from any nonspecific effects of early life anesthesia and surgery, which can potentially induce neuroinflammation and/or neurodevelopmental alterations^40^.

### 2.4 Experiment 1: Acute Neuroimmune Effects of Pediatric Traumatic Brain Injury

The neuroimmune system responds to TBI^19,34–37^ and plays numerous roles in brain development^20,22,28,29^. Thus, we examined markers of microglial and astrocytic coverage, microglial expression of the phagocytic marker CD68, as well as mast cell numbers and degranulation.

#### 2.4.1 Tissue Collection

At 3 days post injury (DPI) [P18], Naïve, Sham, and TBI rats were perfused between 09:00 and 12:00, and brains were collected. Brains were cut into serial 30μm coronal sections and stored in cryoprotectant at -20°C.

#### 2.4.2 IBA1 and GFAP Staining and Analysis

IBA1 (microglia) and GFAP (astrocyte) staining was used to examine glial coverage in HPC, PFC, and Perilesional Cortex^41,42^ (Fig 1D; Table 1). A Zeiss Axioimager M2 microscope and a CX9000 Digital Camera using Steroinvestigator Software (MBF Bioscience) was used to acquire 20x IBA1 and GFAP images on the right side of the brain (ipsilateral to injury) (3-4 images/subregion). In the Perilesional Cortex, images were collected of the retrosplenial (medial to injury) and somatosensory (lateral to injury) corticies (AP-2.0 to -3.0). In the HPC, images were collected of the dentate gyrus (DG), CA1, and CA3 subregions (AP -2.0 to -3.0). In the PFC, images were collected of the infralimbic (IL) and prelimbic (PL) subregions (AP +2.0 to +3.0). ImageJ was used to measure percent area coverage of IBA1 and GFAP. Coverage was analyzed both within subregion and averaged across each region. Image settings were kept constant, and all imaging and analysis was conducted by a blinded observer.

**Table 1:**
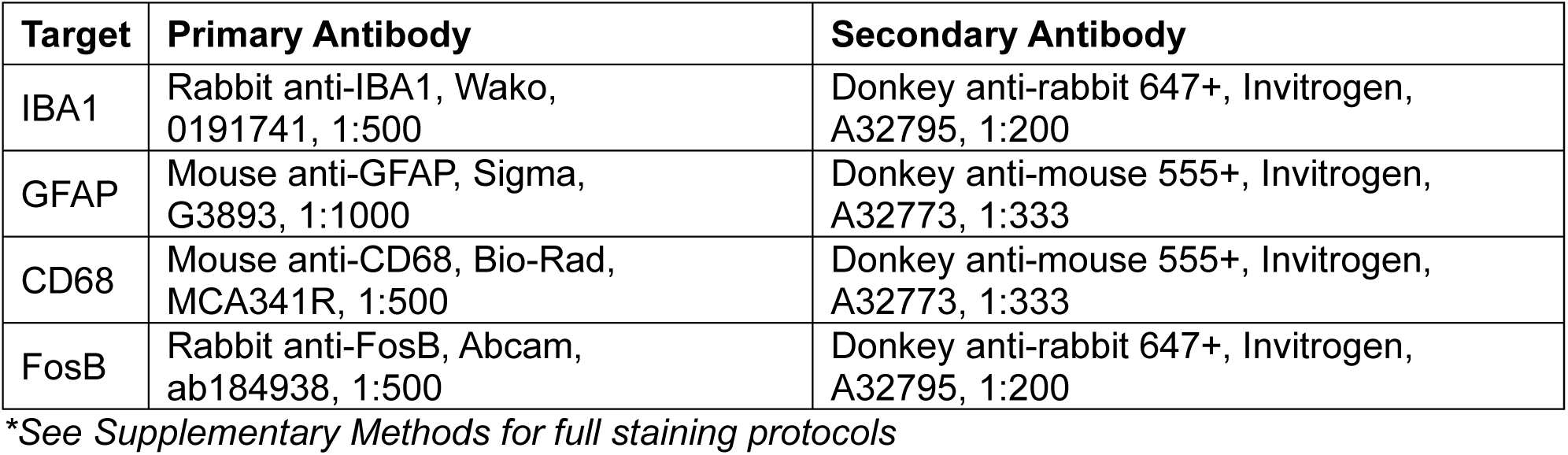
Antibody information.

| **Target** | **Primary Antibody** | **Secondary Antibody** |
| --- | --- | --- |
| IBA1 | Rabbit anti-IBA1, Wako, 0191741, 1:500 | Donkey anti-rabbit 647+, Invitrogen, A32795, 1:200 |
| GFAP | Mouse anti-GFAP, Sigma, G3893, 1:1000 | Donkey anti-mouse 555+, Invitrogen, A32773, 1:333 |
| CD68 | Mouse anti-CD68, Bio-Rad, MCA341R, 1:500 | Donkey anti-mouse 555+, Invitrogen, A32773, 1:333 |
| FosB | Rabbit anti-FosB, Abcam, ab184938, 1:500 | Donkey anti-rabbit 647+, Invitrogen, A32795, 1:200 |

*\*See Supplementary Methods for full staining protocols*

#### 2.4.3 Microglial CD68 Staining and Analysis

Upregulated expression of CD68, a lysosomal marker, in microglia can indicate an increased response to an insult such as TBI^28,43,44^. To assess microglial CD68, a series was co-stained with IBA1 and CD68 (Table 1). Images were collected at 40x using a Zeiss AxioObserver Z.1 with an Apotome digital imaging system and Zen Blue Software (Zeiss). Z-stacks (0.5µm steps) were collected of the same subregions outlined in 2.4.2 (3-4 images/subregion). In Imaris (v10.2.0) the surface tool was used to outline IBA1, and CD68 within this surface was “masked” to assess CD68 expression within microglia. Volumes were collected for IBA1 and masked CD68 surfaces, and normalized CD68 expression within microglia is presented as CD68 volume/IBA1 volume.

#### 2.4.4 Mast Cell Staining and Analysis

Mast cells can be visualized using toluidine blue, which binds to their acidified vacuoles^33,45,45^. A Zeiss Axioimager M2 microscope was used to visualize stained mast cells at 20-40x, and Steroinvestigator was used to tabulate mast cell numbers and degranulation (Fig 4C,D) by a blinded observer. Mast cells were counted across all sections of the HPC (including velum interpositum) and thalamus. Counts for total, granulated, and degranulated mast cells are presented as average number per section.

This stain was also utilized to measure cortical thickness, as part of injury intensity assessment. The distance from the corpus callosum to the outer edge of the cortex was measured at the perilesion site (AP-2.0 to -3.0) for 3 sections per rat in Stereoinvestigator and is presented as average cortical thickness for each animal.

#### 2.4.5 Statistics

TBI and Sex effects were first analyzed by Two-Way ANOVA using GraphPad Prism 10. If Sex or Interaction effects were detected, these results were finalized and are presented below, with males and females shown separately on corresponding graphs. In the absence of Sex or Interaction effects, the data was collapsed across Sex, and a One-Way ANOVA was run to assess main effects of injury. In such cases, graphs show the collapsed sexes together, with different symbols used to denote males (squares) versus females (triangles). Outliers were determined by values that fall outside the mean ± 1.96 times the standard deviation^46,47^. Normality and variance were assessed via Shapiro-Wilk and Brown-Forsythe tests. Tukey’s post hoc tests were conducted when main effects and/or interactions were found. Full statistical results for One-Way and Two-Way ANOVAs are presented in Supplementary Table 1.

### 2.5 Experiment 2: Chronic Behavioral Effects of Pediatric Traumatic Brain Injury

Pediatric TBI exerts lasting behavioral impacts in humans, with alterations to cognitive, social, and mood-related behaviors^2,5,7,11–14^. Thus, we assessed these domains in our rat model (Fig 1D). Behavioral tests were separated by 2-3 days. Unless specified, tests were run between 08:00 and 14:00.

#### 2.5.1 Juvenile Social Play

Rats were weaned into sex- and treatment-matched pairs on P22-24. Juvenile Social Play testing was conducted on P28-P30 [13-15 DPI] at the start of the dark phase under red light after a 4 hr social isolation. Home cage pairs were placed back together in a plexiglass box (19×14.5×12 in) and recorded for 20 min. Play behaviors scored by a blinded observer included chasing, rough and tumble play, and pinning. Data are presented as the total number of play behaviors recorded for each animal across the three sessions.

#### 2.5.2 Spatial Y Maze

On P61 [46 DPI], rats were tested in the Spatial Y Maze under red light. The Y maze has 3 arms (20×5 in each) that stem from center. Spatial cues were placed around and one arm was blocked. Rats were placed in one “starting” arm and allowed to explore for 10 min. After 1 hr, the block was removed and rats were allowed to explore the newly (“novel”) and previously (“familiar”) opened arms for 10 min. Videos were recorded from above. Time spent in each arm and locomotion were quantified using Ethovision 11, and a discrimination index was calculated: (novel duration – familiar duration) / (novel + familiar duration).

#### 2.5.3 Spontaneous Y Maze

Spontaneous Y Maze testing was run on P64 [49 DPI] under red light. The Y maze did not have any arms blocked or spatial cues for this test. Rats were placed in one of the arms and allowed to freely explore for 8 min. Videos were scored by a blinded observer for the sequence of entries into each arm (e.g., “1 to 2 to 3” is spontaneous, “1 to 2 to 1” is not). Data are presented as the percentage of a rat’s entries that are spontaneously alternating.

#### 2.5.4 Social Interaction

On P67 [52 DPI], social interaction was assessed. Rats were placed in an open field (24×24×16 in) under dim white light with a novel age- and sex-matched conspecific for 10 min. Videos of the interactions were scored by a blinded observer. Behaviors quantified included active interaction (e.g., touching, sniffing), passive interaction (e.g., huddling), and avoidance (e.g., moving away from the other rat). Active interaction and avoidance are presented for both experimental and stimulus rats, based on which animal initiated or moved away.

#### 2.5.5 Open Field and Novel Object Recognition

Rats underwent Open Field and Novel Object Recognition Testing on P70-71 [55-56 DPI]. This two-day test was run under dim white light. On day 1, rats were placed in the center of an open field (24×24×16 in) and allowed to explore for 10 min. Two hr later, two objects (500mL bottles filled with blue-dyed water) were placed in the open field, and rats were allowed to explore for 10 min. The next morning, rats were placed back in with the same 2 objects for 10 min. Four hr later, one of the objects was changed to a 5×4×3 in pink tube rack, and rats were allowed to explore both objects for 10 min. Ethovision 11 was used to assess center duration and locomotion during the first phase (Open Field) and time spent exploring the novel and familiar objects during the last phase (Novel Object Recognition). A discrimination index was calculated: (novel duration – familiar duration) / (novel+familiar duration). A more positive index indicates a higher preference for the novel object.

#### 2.5.6 Elevated Plus Maze

On P76 [61 DPI] rats were tested in an Elevated Plus Maze under red light. This maze consists of a center area (5 in^2^), two open arms (20×5 in) and two closed arms (20×5×15 in). Rats were placed on one of the open arms, facing center, and allowed to explore for 5 min. Videos were recorded from above and Ethovision 11 was used to assess time spent in the open arms, closed arms, and center, as well as locomotion.

#### 2.5.7 Statistics

Statistical testing largely matched that described in 2.4.5. Juvenile Social Play data was analyzed as a Two-Way Mixed ANOVA (TBI x Day).

### 2.6 Experiment 3: Risky Decision-Making Effects of Pediatric Traumatic Brain Injury

The Rodent Gambling Task (RGT) is the rodent version of the Iowa Gambling Task, yielding information about risky decision-making and impulsivity^48,49^ (Fig 1D). Here, we used the RGT to gain insight into potential deficits in these behavioral domains after pediatric TBI.

#### 2.6.1 Juvenile Social Play and Spontaneous Y Maze

A new cohort of rats was generated. Prior to the RGT, they were tested in Juvenile Social Play (P28-30) and the Spontaneous Y Maze (P64) as described above, to increase statistical power for those datasets. Results were combined with the corresponding Experiment 2 results for analysis and presentation.

#### 2.6.2 Housing and Training

On P65, rats were transferred to a new housing building for access to operant boxes, switched to an inverted light cycle, and food restricted to 85% of ad libitum body weight. Operant testing consisted of daily 30 min sessions (Monday-Friday)^50,51^. From P71-91 [56-76 DPI], rats were habituated to the boxes and trained to nose poke a lit hole for a sugar pellet reward. In the latter phase of training, they underwent a week of forced choice, which familiarized them with the win/loss rules of Choices 1-4 (described below). An additional requirement to withhold responding for 5 s prior to choice served as a measure of impulsivity.

#### 2.6.3 Rodent Gambling Task Procedure

Following training, rats entered free choice RGT, in which they had a choice between 4 nose poke holes (Fig 6A). Each hole had different probabilities and magnitudes of reward (# sugar pellets) or punishment (timeout with lights on). Choice 2 represented the optimal choice to receive the most pellets. Choice 1 was safer yet suboptimal, yielding consistent but small rewards. Choices 3 and 4 represented gradually more risky decisions, offering more reward but longer punishment. Rats tested for 30 sessions (6 weeks). Custom Med-PC IV software was used to record and export the results of each trial. Data were aggregated into 6 5-session bins for analysis and presentation.

The primary endpoint of this task is percent choice for each option (Choices 1-4). Overall performance across all 4 choices can also be represented by an RGT Score metric. Score demonstrates the shift from the safer Choices 1 and 2 to the riskier Choices 3 and 4, with higher scores representing more optimal performance. The secondary measure of this task is percent of premature responses (i.e. nose poking prior to the hole lighting up), which offers insight into impulsivity and disinhibition. Another endpoint of interest in this cohort is WinStay (i.e. sticking to a previous choice that was rewarded), which offers insight into task optimization. Additional variables the RGT yields are Omissions, Number of Trials, Pellets Earned, Choice Latency, Stay, and LoseStay (see Supplementary Methods).

#### 2.6.4 Tissue Collection and FosB Assessment

Following completion of the RGT, rats were sacrificed and brains were collected. One series was stained for FosB analysis in regions previously implicated in the RGT^51^, namely the PFC [IL and PL] and nucleus accumbens (NAc) [Shell]. A FosB stain was run (Table 1), and images were acquired as described in 2.4.2. The ImageJ counting tool was used by a blinded experimenter to quantify the number of FosB+ cells.

#### 2.6.5 Statistics

Statistical testing was performed in R (v4.3.4) using the packages *lmerTest, MuMIn, brms,* and *emmeans*. Tests were run on both acquisition (all sessions) and stable (last 5 sessions) data. Given that choice data is interdependent (e.g. making one choice sacrifices the ability to make the others), Bayesian models with choice as a random effect were used to assess TBI-related decision-making effects. Results were considered significant by credible intervals that did not cross zero. Linear mixed models were used to assess Score, Premature, and WinStay data, as well as the additional RGT variables outlined above. Estimated marginal means were used for post-hoc comparisons. FosB results were analyzed using GraphPad Prism 10 via Two-Way and One-Way ANOVAs as described in section 2.4.5.

#### 2.6.6 Exploratory Analyses

Since there was not sufficient power for full sex difference analyses, exploratory analyses were conducted to investigate the potential role of sex in RGT endpoints. Bayesian models containing sex as an interaction term (Sex*Injury) or main effect (Sex+Injury) were compared to the Injury alone model using Bayes factors for acquisition and stable choice data. Linear mixed models were run with sex as factor for other variables that exhibited injury effects. Due to conceptual similarities between WinStay and Y Maze results, Pearson correlation analyses between these endpoints were run for each sex using GraphPad Prism 10.

## RESULTS

### 3.1 TBI Intensity Assessment

The present 2atm TBI elicited righting times of ∼3-5min (Fig 1B), demonstrating that the injury was stronger than in our previous study^40^ using a 1.2 atm LFPI, which elicited ∼1min righting times. We also observed no signs of tissue cavitation or changes in cortical thickness near the site of injury at 3DPI (Fig 1C; F_Injury_(2,32)=0.002, p=0.998). These measures together suggest a mild injury model^52,53^.

### 3.2 Experiment 1: Acute Neuroimmune Effects of Pediatric Traumatic Brain Injury

#### 3.2.1 TBI increased IBA1 and GFAP in the Perilesional Cortex but not the HPC or PFC

At 3DPI, TBI significantly increased IBA1 percent area in the Perilesional Cortex (Fig 2A-B; F_Injury_(2,32)=10.87, p<0.001), relative to both Naïve (p<0.001) and Sham (p=0.024). No injury effects on IBA1 staining were seen in the HPC (Fig 2C; F_Injury_(2,32)=2.047, p=0.146) or PFC (Fig 2D; F_Injury_(2,32)=0.698, p=0.505). GFAP exhibited a similar pattern, with TBI significantly increasing GFAP percent area in the Perilesional Cortex (Fig 2E-F; F_Injury_(2,32)=10.42, p<0.001) relative to Naïve (p<0.001) and Sham (p=0.002), but not the HPC (Fig 2G; F_Injury_(2,31)=0.030, p=0.971) or PFC (Fig 2H; F_Injury_(2,31)=0.204, p=0.816). These overall patterns persisted in respective subregions (Supplementary Fig 1A-B). No sex differences were observed in IBA1 (F_Sex_(1,29)=0.118, p=0.734) or GFAP (F_Sex_(1,29)=0.113, p=0.739). These results suggest that pediatric TBI increases glial coverage locally but not globally.

**Figure 2:**
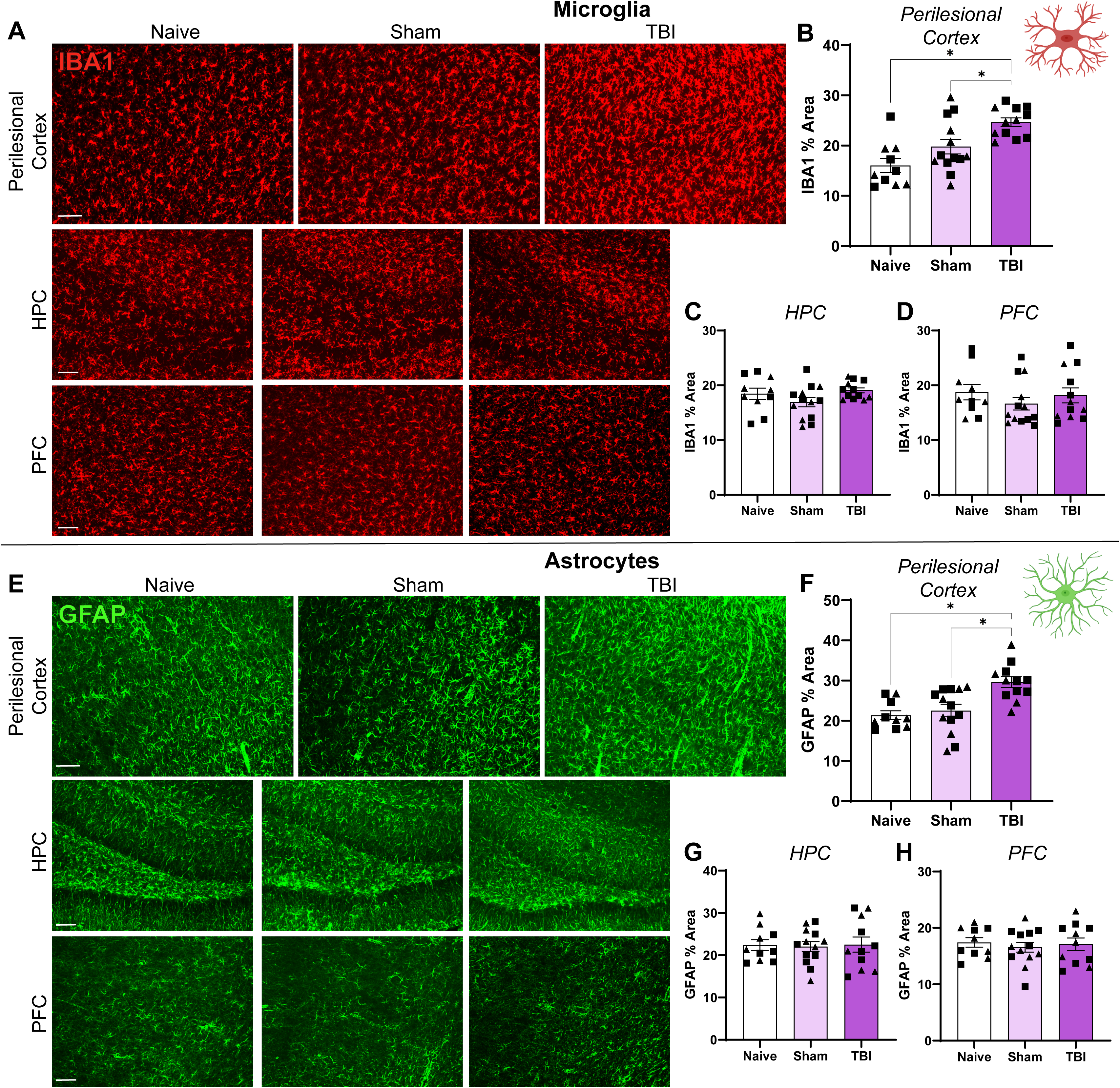
Acute local increases in glial coverage following pediatric TBI. (A) IBA1 representative images (scale bar = 100µm). (B-D) TBI increased IBA1 percent area in the Perilesional Cortex, but not the HPC or PFC. (E) GFAP representative images (scale bar = 100µm). (F-H) TBI increased GFAP percent area in the Perilesional Cortex, but not the HPC or PFC. *p<0.05. N=5-7/sex /group. Error bars= Mean±SEM. Males are denoted by squares, while females are denoted by triangles.

#### 3.2.2 TBI increased microglial CD68 in the Perilesional Cortex, HPC, and PFC

TBI increased the volume of microglial CD68 throughout the regions analyzed. In the Perilesional Cortex, TBI increased in CD68 volume (Fig 3A-B; F_Injury_(2,32)=45.56, p<0.001) relative to both Naïve (p<0.001) and Sham (p<0.001) in both sexes. In the HPC, TBI also increased CD68 volume (Fig 3C; F_Injury_(2,27)=19.50, p<0.001), but a sex x injury interaction (F_Interaction_(2,27)=4.941, p=0.015) emerged. TBI females exhibited a significant increase in CD68 volume (p<0.001 vs Naïve and Sham), but the increase in males did not reach significance (p>0.10 vs Naïve and Sham), and females’ TBI CD68 volume was higher than males’ (p<0.001). In the PFC, TBI caused a subregion-specific increase in CD68 volume. While no differences were observed in the overall PFC or PL (Supplementary Fig 2), CD68 volume was significantly increased in the IL in both sexes (Fig 3D; F_Injury_(2,27)=4.918, p=0.015) relative to Naïve (p=0.034) and Sham (p=0.029). While there is nuance by sex and subregion, this increase in microglial CD68 suggests that pediatric TBI causes microglial activation distal from the site of injury in the absence of changes in glial coverage.

**Figure 3:**
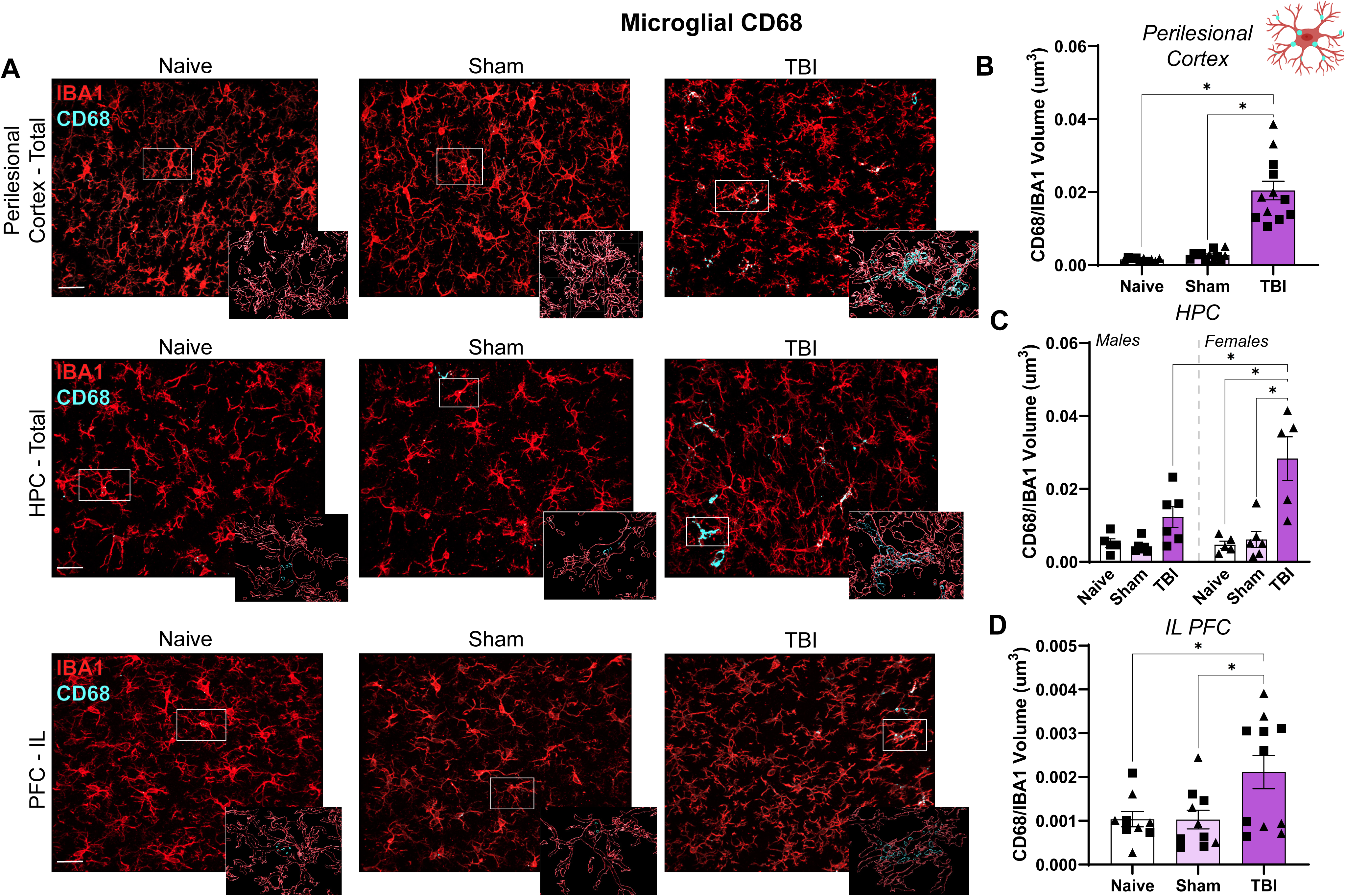
Acute widespread increases in microglial CD68 following pediatric TBI. (A) Representative images (scale bar = 30 µm) and Imaris reconstructions of CD68 expression within microglia. (B) TBI greatly increased CD68 in the Perilesional Cortex. (C) TBI increased CD68 in the HPC, to a higher degree in females than males. (D) TBI increased CD68 specifically in the infralimbic (IL) PFC. *p<0.05. N=4-7/sex/group. Error bars= Mean±SEM. Males are denoted by squares, while females are denoted by triangles.

#### 3.2.3 TBI increased mast cell number and degranulation

TBI increased the number of mast cells throughout the HPC, velum interpositum, and thalamus (Fig 4A-B; Supplementary Fig 3; F_Injury_(2,32)=5.943, p=0.006) relative to both Naïve (p=0.047) and Sham (p=0.007). This increase was largely driven by degranulated cells (Fig 4C-F; F_Injury_(2,32)=8.148, p=0.001), indicating that mast cells are not only more abundant but also more activated acutely following TBI (p=0.007 vs Naïve; p=0.002 vs Sham). No sex differences were observed in mast cell number (F_Sex_(1,29)=0.722, p=0.403) or degranulation (F_Sex_(1,29)=0.430, p=0.517).

**Figure 4:**
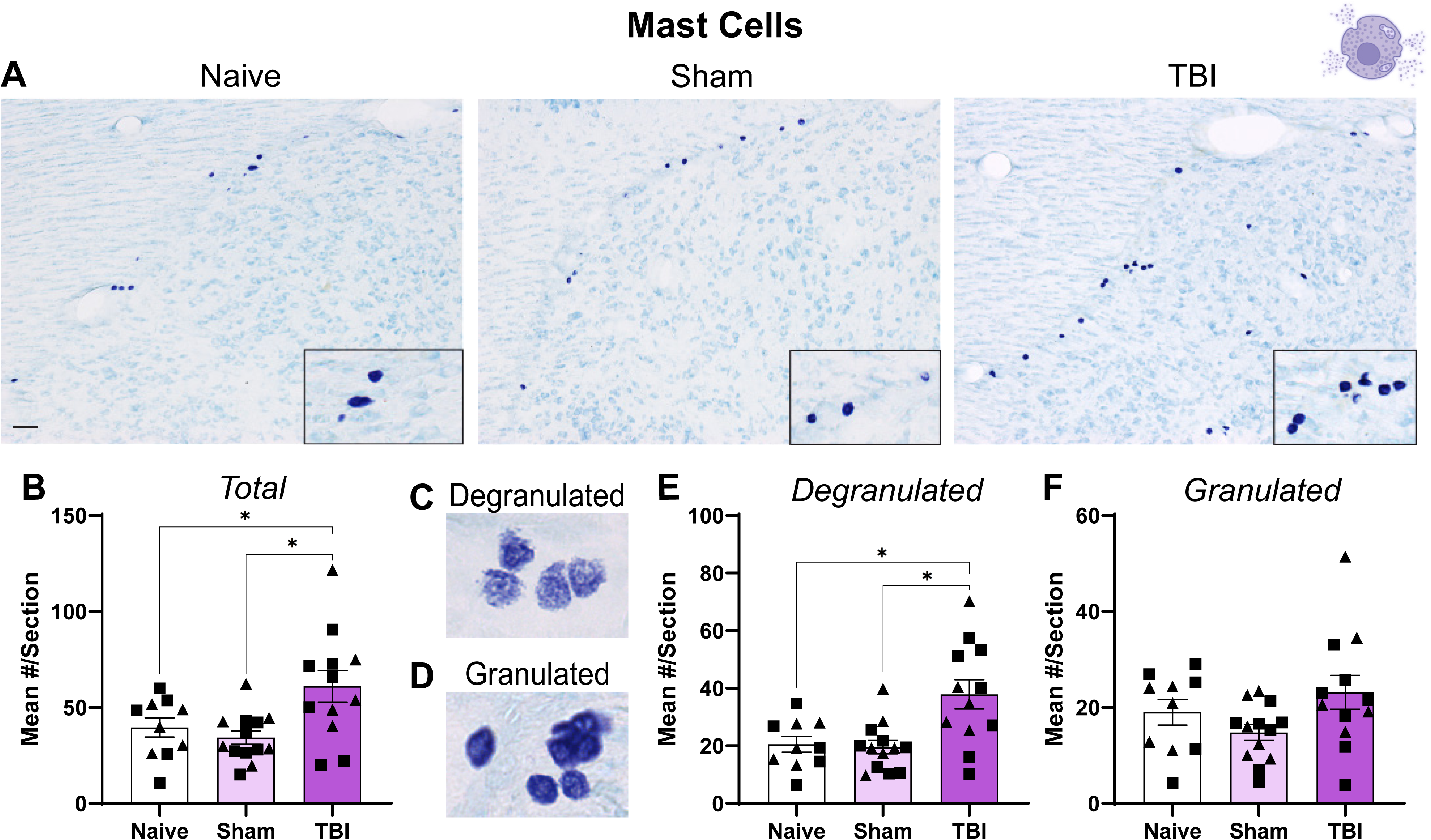
Acute mast cell activation following pediatric TBI. (A) Representative images of mast cells (dark purple) in the velum interpositum underneath the HPC (scale bar = 100 µm). (B) TBI increased total mast cell numbers throughout the brain. Mast cells were further characterized as degranulated (diffuse; more active state) or granulated (dense; less active state). (C-D) Representative images of degranulated vs granulated mast cells. (E-F) The TBI-induced increase in mast cells was driven by degranulated, rather than granulated mast cells. *p<0.05. N=5-7/sex /group. Error bars= Mean±SEM. Males are denoted by squares, while females are denoted by triangles.

### 3.3 Experiment 2: Chronic Behavioral Effects of Pediatric Traumatic Brain Injury

#### 3.3.1 TBI altered cognitive performance in adulthood

We did not observe TBI effects in the Spatial Y Maze (Fig 5A; F_Injury_(2,18)=0.810, p=0.461). There was an effect of sex (F_Sex_(1,18)=5.211, p=0.035) and a sex x injury interaction (F_Interaction_(2,18)=3.920, p=0.039), in that females exhibited higher discrimination than males in Naïve (p=0.028) and Sham (p=0.012), but not TBI (p=0.345), groups. No post hoc tests were significant between injury groups (p’s>0.058), so spatial memory does not appear to be affected by TBI in this task. Distance traveled was not affected by injury (Supplementary Fig 4A; F_Injury_(2,21)=0.736, p=0.491) or sex (F_Sex_(1,18)=3.388, p=0.082).

**Figure 5:**
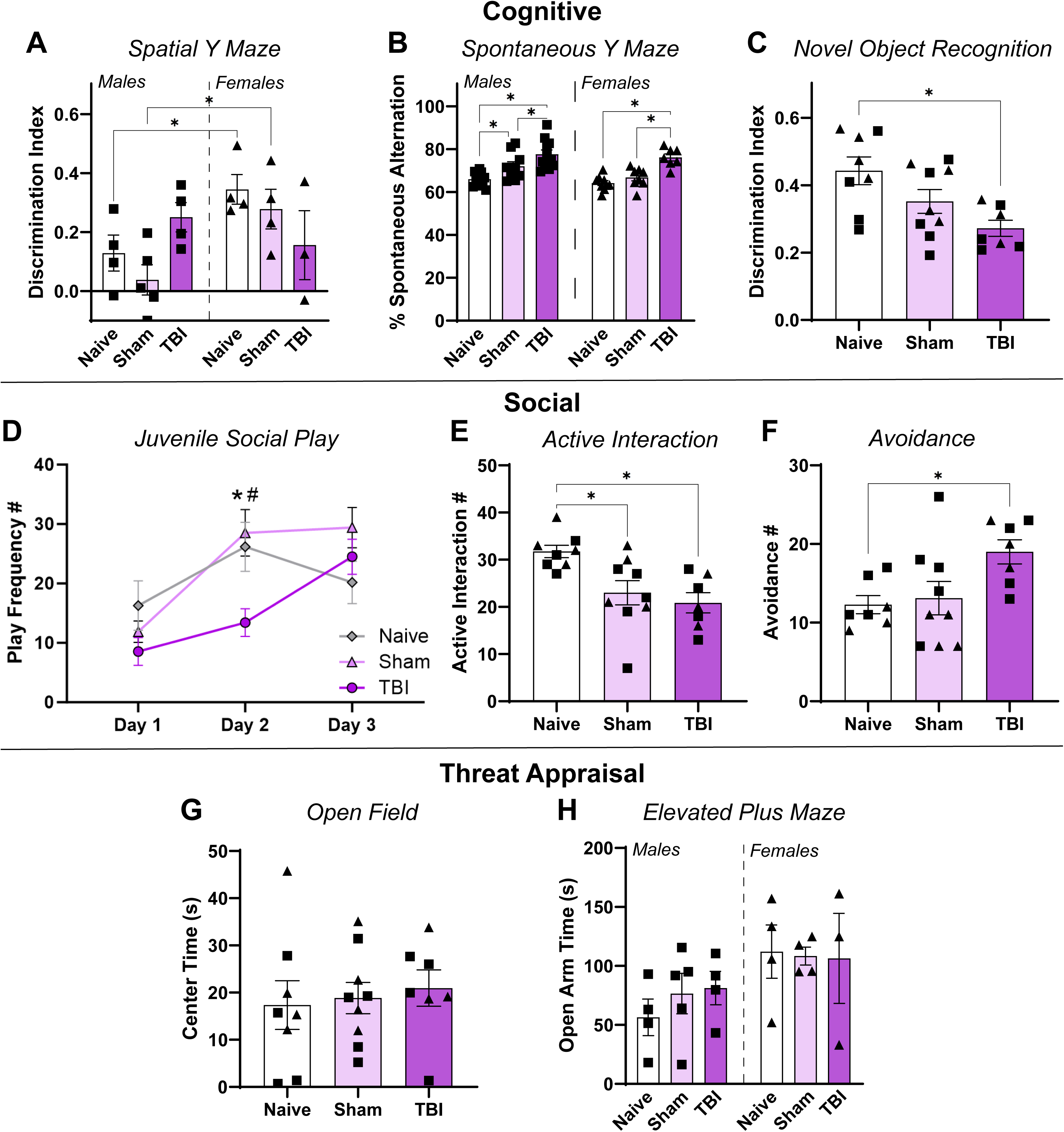
Pediatric TBI impacted cognitive and social behavior. Cognitive behaviors: (A) TBI did not impact performance in the Spatial Y Maze, although females did show higher discrimination than males on this task in general. (B) TBI increased spontaneous alternation in the Spontaneous Y Maze, with males showing higher spontaneity than females. (C) TBI impaired discrimination in the Novel Object Recognition task in both sexes. Social behaviors: (D) TBI reduced juvenile social play in both sexes, primarily on day 2 of testing. * = TBI vs Naïve. ^#^ = TBI vs Sham. (E-F) TBI and Sham reduced active social interaction, while TBI alone increased social avoidance in both sexes. Mood-related behaviors: (G-H) TBI did not impact performance in the Open Field or Elevated Plus Maze. Females did spend more time in the open arms than males in general. *p<0.05. N=3-5/sex/group for A, C, and E-H. N=8-11/sex/group for B and D. Error bars= Mean±SEM. Males are denoted by squares, while females are denoted by triangles.

For the Spontaneous Y Maze, TBI rats exhibited a higher rate of spontaneous alternation (Fig 5B; F_Injury_(2,48)=24.57, p<0.001) than Naïve (p<0.001) and Sham (p<0.048) rats. There was also a sex difference (F_Sex_(1,48)=4.053, p=0.0497); however, this was likely an artifact driven by females’ higher entries during the task (F_Sex_(1,18)=13.45, p<0.001).

TBI impaired Novel Object Recognition. While all groups discriminated the novel object (higher than zero), the discrimination index was significantly lower in TBI rats than Naïve rats (Fig 5C; F_Injury_(2,21)=5.581, p=0.011; TBI vs Naïve p=0.009). Sham rats fell in the middle but did not significantly differ from Naïve (p=0.166) or TBI (p=0.268). These results suggest that TBI impairs, but does not eliminate, memory performance in this task. Sex did not affect discrimination (F_Sex_(1,18)=3.335, p=0.084). Females had a higher distance traveled than males (F_Sex_(1,18)=8.159, p=0.011), but injury did not impact locomotion (F_Injury_(2,18)=0.073, p=0.930).

#### 3.3.2 TBI impaired juvenile and adult social behavior

Juvenile social play was assessed between sex- and treatment-matched cage mates for 3 sessions. There was a significant interaction between testing day and injury (Fig 5D; F_Interaction_(4,96)=5.161, p<0.001)), with TBI rats exhibiting less play than Naïve or Sham rats, primarily on day 2 (p=0.029 vs Naïve, p=0.003 vs Sham). These results suggest that TBI may delay habituation to the testing arena or enhance the salience of social deprivation prior to testing, since TBI rats did not show the increase in play across sessions that Naïve and Sham rats showed on day 2 until day 3.

In adulthood, we examined social interaction with a sex-matched novel conspecific. TBI and Sham both decreased active social interaction (Fig 5E; F_Injury_(2,21)=7.105, p=0.004; TBI vs Naïve p=0.006, Sham vs Naïve p=0.019). TBI alone increased avoidance (Fig 5F; F_Injury_(2,20)=4.016, p=0.034; TBI vs Naive p=0.047, Sham vs Naive p=0.067). There was no difference in passive interaction (Supplementary Fig 4B; F_Injury_(2.21=0.058, p=0.943). Avoidance of the experimental rat by the stimulus rat did show an injury effect (F_Injury_(2.18)=4.646, p=0.024); although no post hoc tests were significant (p’s>0.09).

#### 3.3.3 TBI did not impact threat appraisal

In the Open Field, TBI did not affect center time (Fig 5G; F_Injury_(2,21)=0.176, p=0.840). Females had a higher distance traveled than males (F(1,18)_Sex_=26.18, p<0.001), but there were no effects of injury (F_Injury_(2,18)=0.086, p=0.918) or sex x injury interactions (F_Interaction_(2,18)=0.086, p=0.918) on locomotion.

A similar pattern occurred in the Elevated Plus Maze, with no effect of TBI on open arm time (Fig 5H; F_Injury_(2,18)=0.140, p=0.870). There was a sex effect (F_Sex_(1,18)=5.570, p=0.030), with females spending more time in the open arms; although, post hocs were not significant (p’s>0.056), and this could be because females had a higher distance traveled than males (F_Sex_(1,18)=6.351, p=0.021).

### 3.4 Experiment 3: Risky Decision-Making Effects of Pediatric Traumatic Brain Injury

#### 3.4.1 TBI increased risky decision-making

The main endpoint of the RGT is choice: Choice 2 is optimal, Choice 1 is suboptimal, and Choices 3 and 4 are increasingly risky (Fig 6A). Naïve rats quickly showed a preference for the optimal Choice 2 (Fig 6C; Supplementary Table 1). While TBI rats still picked Choice 2 the most, they exhibited a ∼20% shift towards the riskier Choice 3 (Fig 6D). Sham rats fell between Naïve and TBI rats. Bayesian analyses revealed a significant injury x session interaction (CI - 0.129 to -0.056) over the entire curve (“acquisition”), as well as a significant injury effect (CI - 1.554 to -0.231) in the last testing week (“stable”). Post hocs indicate that TBI rats made Choice 2 significantly less (CI 1.273 to 0.118) and Choice 3 significantly more (CI -0.061 to - 1.522) than Naïve rats, while Sham rats did not significantly differ from either Naïve or TBI. Choices 1 and 4 did not differ by injury condition (Fig 6B,E). In terms of overall performance, TBI significantly decreased RGT Score (Fig 6F). There was a significant injury x week interaction (F_Interaction_(2, 867)=18.67; p<0.0001), with TBI rats developing less preference for safe options than Naïve and Sham rats (p’s<0.001), reinforcing the above finding of increased risky decision-making.

**Figure 6:**
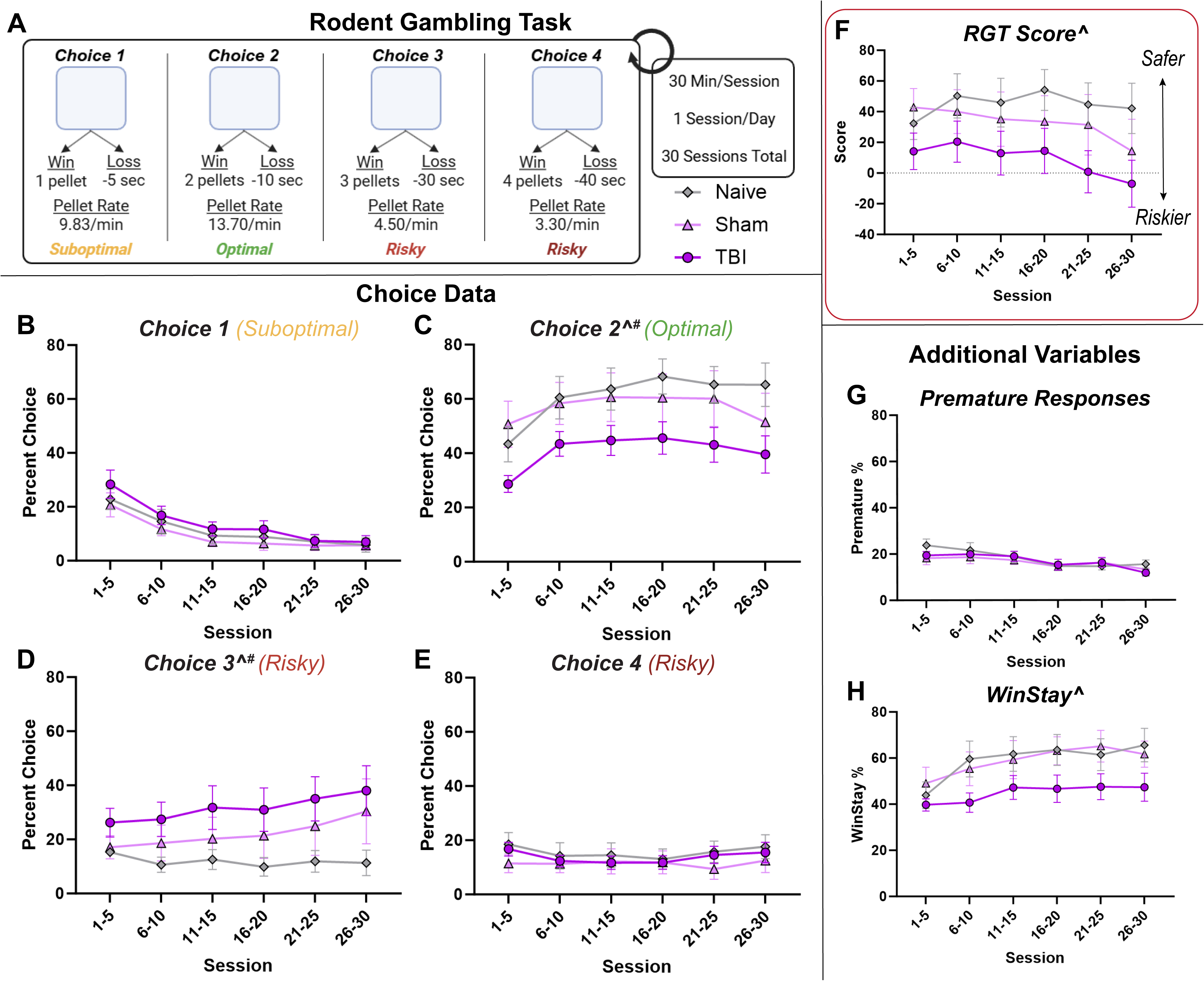
Pediatric TBI increased risky decision-making in adulthood. (A) Rodent Gambling Task design and win/loss information for Choices 1-4, which range from safe to risky. Choice 2 yields the optimal reward (pellet rate). Rats are trained on these conditions then undergo daily testing for 30 sessions. (B-E) Percent choice curves across sessions. While TBI did not alter Choice 1 or Choice 4, it did lead to a shift from Choice 2 to Choice 3, suggesting more risky decision-making. (F) TBI also reduced RGT score, indicating more risky decisions overall. (G, H) Additional RGT variables. TBI did not impact premature responding but did reduce WinStay percentage. ^ = TBI x Time Interaction (“acquisition”). # = Main effect of TBI (“stable”). N=4-7/sex/group. Error bars= Mean±SEM.

#### 3.4.2 TBI did not impact impulsivity but reduced task optimization

TBI did not impact premature responding (Fig 6G; F_Interaction_(2, 867)=2.346; p=0.096), suggesting that this model does not affect impulsivity. In contrast, there was a significant injury x week interaction for the WinStay measure (Fig 6H; F_Interaction_(2, 867)=3.821; p=0.022), with TBI rats exhibiting decreased WinStay behavior relative to Naïve rats (p=0.018). This shift suggests that TBI rats switched their choices more frequently even when they were being rewarded, which is indicative of impaired task optimization. TBI did not impact other endpoints of interest (Supplementary Table 1), including Omissions and Choice Latency (i.e., inattention), Number of Trials and Pellets Earned (i.e., overall performance), or Stay and LoseStay (i.e., perseveration). This suggests that this pediatric TBI model primarily impacts risky decision-making, and possibly behavioral optimization, rather than these other cognitive domains.

#### 3.4.3 FosB was not impacted by TBI

There were no effects of TBI on the number of FosB+ cells in the IL (Supplementary Fig 5F; F_Injury_(2,26)=1.877, p=0.173), PL (F_Injury_(2,26)=1.940, p=0.164), or NAc (F_Injury_(2,25)=2.062, p=0.148). No sex differences were observed in FosB expression (p’s>0.451).

#### 3.4.4 Exploratory Analyses: Sex Effects and RGT-Y Maze Relationships

While underpowered, visually there appeared to be sex differences in RGT performance, with a potential main effect of females better optimizing for Choice 2 or an interaction of sex with injury (Supplementary Fig 5A-B). Indeed, comparative analyses revealed that accounting for sex significantly improved model fit. For acquisition data, including sex as a main effect (Injury + Sex) provided the strongest model fit (7.69 log Bayes factor difference vs Injury), suggesting that sex influenced a rat’s performance throughout the RGT, regardless of injury. For stable data, including sex as an interaction term (Injury*Sex) provided the strongest model fit (2.31 log Bayes factor vs Injury), suggesting that both sex and injury determine where a rat’s choice ultimately settled. Sex also affected RGT Score (Supplementary Fig 5C; Sex x Injury x Week: F_Interaction_(2, 864)=4.44; p=0.012) and WinStay (Supplementary Fig 5D; Sex x Injury: F_Interaction_(1, 864)=3.41; p=0.047), with females driving the TBI-induced reduction in these endpoints.

Furthermore, WinStay behavior was negatively correlated with Y maze spontaneous alternation (Supplementary Fig 5E), such that rats that were less likely to stay with a rewarded choice were also more likely to be more spontaneous in the Y maze. Importantly, this correlation was only present in females (R^2^=0.307, p=0.049), not in males (R^2^=0.007, p=0.774), further suggesting sex differences in this switching phenotype.

## DISCUSSION

Here, we demonstrated that a mild (2atm) TBI delivered during the late neonatal period (P15) in rats induced lasting behavioral effects reminiscent of those seen in humans, and acute neuroinflammation may be an intriguing candidate underlying these effects. At 3DPI, pediatric TBI precipitated local increases in glial coverage, along with widespread increases in microglial CD68 expression and mast cell activation. In juvenile and early adult rats, TBI altered cognitive and social function, but not anxiety-related behavior. Later in adulthood, TBI increased risky decision-making and impaired task optimization, especially in females. Together, these results provide compelling face validity and set the stage for future studies aimed at understanding the mechanistic link between acute neuroinflammation during critical periods of development with prolonged behavioral impairment following pediatric TBI (Fig 7).

**Figure 7:**
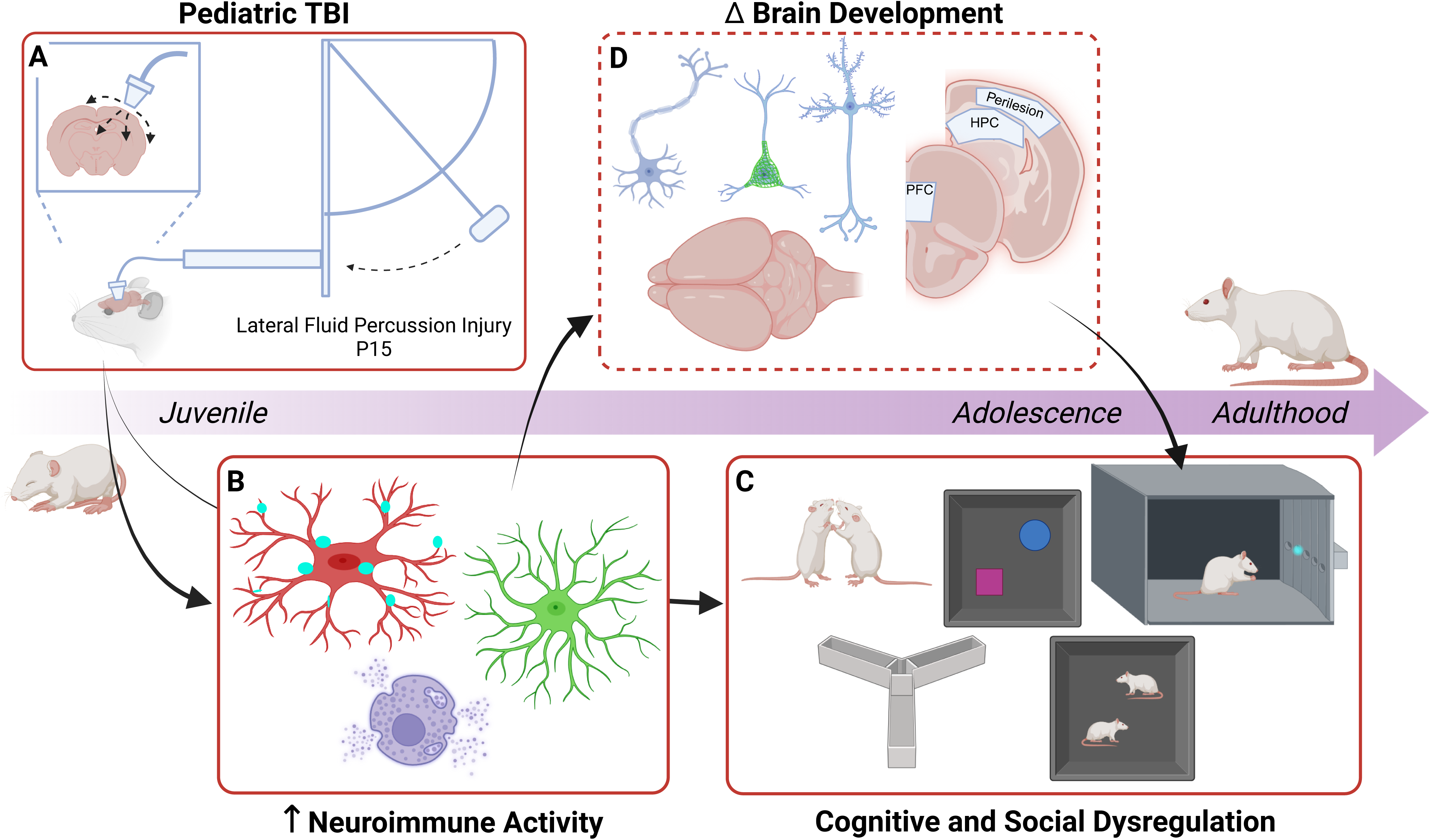
Summary Figure. (A) A mild TBI experienced at P15 in male and female rats causes (B) acute activation of microglia, astrocytes, and mast cells, as well as (C) long-lasting cognitive and social dysregulation. These effects could be linked, in that (D) neuroinflammation during this critical period stands to alter how neurons develop in key behavior-regulating brain regions.

While our prior study with a milder injury showed some social impairments^40^, the present model induced a greater number of more robust cognitive and social phenotypes. Rats that experienced pediatric TBI showed reduced novel object recognition and active social interaction, along with increased risky decision-making and social avoidance. The existing literature on pediatric TBI is heterogenous^27,38,54^, featuring different models (LFPI, medial FPI, weight drop, controlled cortical impact (CCI), repeated CCI), ages (P7-46), intensities (mild-severe), and species (rats vs mice), yet the present results generally align with prior behavioral findings. Studies have identified acute and chronic deficits in memory^55–57^, sociability^58–60^, and even decision-making^61–63^. Some have identified mood-related dysfunction^56,64^; although, we did not observe anxiety-related behavior here. Mood-related changes may be more prominent following more severe CCI injuries^56,64^ or when pediatric TBI is experienced in conjunction with additional stressors^65,66^. Digging deeper into cognitive effects, the RGT is a long-term operant task that elicits stable, trait-level behaviors analogous to those seen in humans and allows for dissociation of impulsive responding from risky decision-making^51,67^. The lack of impulsive responding in our pediatric TBI rats suggests that the risky phenotypes seen here are indicative of shifts in decision-making, rather than alterations to disinhibitory control. Interestingly, pediatric TBI rats demonstrated decreased WinStay behavior, along with increased spontaneous alternation in the Y maze, both of which suggest enhanced behavioral switching. Switching refers to an inability to stick with one option, leading to a failure to optimize which could contribute to poorer overall decision-making^49,68^. Overall, the behavioral phenotypes seen here largely reflect those seen in children who experienced a TBI, who often display impaired executive function and social withdrawal both in and beyond school age^5,8,10,11^, positioning it as a translationally-relevant model to interrogate underlying mechanisms in future studies.

These behavioral effects extending through different developmental stages supports a hypothesis in which acute perturbations (such as TBI) during critical periods rewire key circuits, inducing lasting differential connectivity and function, which impacts behavior throughout that animal’s life^21,22,38,39^. To that end, we sought to identify potential candidates that play important roles in both brain development and TBI. As the neuroimmune system fit both qualifications^28,29,34,36^, we investigated the acute impact of pediatric TBI on microglia, astrocytes, and mast cells. At 3DPI, we observed increased coverage of microglia and astrocytes near the site of injury, but not extending to the HPC or PFC. Despite this lack of gross morphological change, microglia in the HPC and IL PFC exhibited increased CD68 expression, suggesting that microglia are more phagocytic^43,44^, and thus likely more engaged in functional changes after TBI compared to microglia in Naïve and Sham animals. These findings align with prior studies demonstrating increased microglia coverage^64,69,70^, astrocyte coverage^64,71^, and CD68 expression^64,69,72^ following various forms of pediatric TBI. Mast cells also increased in both number and degranulation state, indicating another point of heightened immune activation following pediatric TBI. Importantly, these changes were restricted to TBI animals, not animals that received Sham surgery, suggesting that the injury itself played a key role in this neuroinflammation.

This heightened neuroinflammation could impact the typical neurodevelopmental roles of these immune cells. For example, microglia prune developing synapses to shape connectivity, release cytokines to influence neurons and other glia via signaling alterations, and remodel the extracellular matrix to promote or limit neuroplasticity^22,28,29^. If, as our data suggests, microglia are more active during postnatal neurodevelopmental critical periods as a result of TBI, these mechanisms could become aberrant and change how key circuits are forming, as has been demonstrated in early life stress studies^73,74^. While pediatric TBI studies have not directly assessed synaptic pruning, some have used Golgi-Cox staining to assess dendritic morphology. Studies have found increased^61^, decreased^59,63^, or sex-dependent changes^59,63^ in dendritic complexity, likely again due to differences in injury, age, and region. On a broader level, a few studies examining microglia inhibition acutely following CCI have demonstrated worse outcomes on spatial memory, attributing these deficits to a reduced capacity to clear dying neurons after injury^70,75^. These studies demonstrate the potential complexity of the role(s) microglia play in pediatric TBI and highlight the importance of better understanding the molecular underpinnings of their various functions. Future studies should focus on characterizing the molecular landscape of microglia acutely following pediatric TBI as well as their broader impact on neurodevelopment. Similarly, alterations to astrocytes and mast cells could also impact development, via direct interaction, signaling molecules, effects on microglia, etc.^31,76^, and should continue to be considered in future studies.

One interesting note is the limited sex differences observed in the present studies. While sex differences after pediatric TBI are noted in humans, the most notable differences relate to enhanced aggression in boys and more prevalent depression and anxiety symptoms in girls^11,15–17^. We did not directly assess aggressive or depression-like behaviors here, so it possible that different tests, such as a tube test or forced swim test, would reveal more sex differences in behavioral domains of relevance to human deficits. Indeed, the cognitive and social effects seen here do tend to appear in both men and women^12,17,39^. As suggested above, perhaps a stronger injury or additional insult would reveal more sex differences in our anxiety-related endpoints. One exception is the RGT, which did reveal stronger phenotypes in females than males. The RGT is a rodent analog of the Iowa Gambling Task^48,49^, which can elicit worse performance in women who previously experienced a pediatric TBI^16^, making a potential sex difference in this endpoint translationally relevant.

The neuroimmune system also shows sex differences during development that did not seem to be impacted here. While females did exhibit more HPC CD68 than males, the acute immune response to TBI was largely seen in both sexes. It is possible that the TBI was strong enough to supersede underlying sex differences. Our prior study with a weaker TBI did suggest a sex difference in mast cells^40^ that was not present with the slightly stronger injury here. It could also be the case that we are simply at the wrong developmental timepoint to observe sex differences. For example, microglia numbers normally decrease in the HPC earlier in males than females^22,28,29^, so assessing later timepoints would offer greater insight into the dynamics of their response in males versus females. As a caveat, these studies may have been underpowered to detect more mild sex differences. Given that some suggestion of sex differences appeared in our dataset, future studies will include higher replicate numbers to better assess for sex differences.

Another consideration in future studies will be the impact of outside environmental factors. Children do not experience TBI in vacuum, and life history can influence pediatric TBI risk, treatment, and outcomes. There is clinical evidence that poverty, parental absence, and abuse can worsen TBI symptoms later in life^18,65,66,77^. Thus, examining the interaction between factors such as early life stress or environmental enrichment with pediatric TBI could better contextualize our understanding of the role of the neuroimmune system in the developing brain, and shed light on how the complexities of lived experiences influence broader microglia function.

## CONCLUSIONS

Pediatric TBI can lead to struggles throughout a person’s life, yet how this early life challenge exerts such lasting impacts remains largely unknown. Here, we demonstrated that a mild LFPI delivered at P15 in male and female rats 1) induces acute neuroinflammatory profiles during a developmental critical period and 2) generates multiple cognitive, social, and decision-making phenotypes similar to those seen in humans. These findings provide face validity for the present model and set the stage for mechanistic inquiries into the relationship between aberrant activation of the neuroimmune system, altered developmental trajectories of key regions and circuits, and lasting behavioral dysregulation following pediatric TBI. Future studies will seek to identify specific molecular candidates that could be targeted to ameliorate the chronic symptoms of pediatric TBI and improve quality of life for children who undergo this early challenge.

### Transparency, Rigor, and Reproducibility Summary

This study was not formally registered because it was an exploratory basic study designed to establish a new model. The analysis was not formally pre-registered, but the first author certifies that the analysis plan was pre-specified. A sample size of 8-12 rats per group, with groups composed equally of male and female rats, was planned based on prior histochemical and behavioral studies from our lab (to achieve 80% power to detect significant main effects and interactions with medium-large effect sizes (η2p = 0.3-0.8 on one-way and two-way ANOVAs; Cohen’s d= 0.7 on post hocs)). Final N’s are as follows: Experiment 1 N = 35 (10-13 rats/group; 5-7 rats/sex/group), Experiment 2 N = 26 (8-9 rats/group; 4-5 rats/sex/group), Experiment 3 N = 31 (8-12 rats/group; 4-7 rats/sex/group). In Experiment 2, 2 TBI females died between juvenile social play testing and adult behavioral testing as a result of housing complications. Subjects were randomly assigned to groups on the day of TBI. Randomization was stratified by sex and litter, such that 1-3 pups/sex/litter were represented in each group. Analyses of experimental endpoints were performed by investigators blinded to relevant characteristics of the subjects. Experimental manipulations (TBI, behavioral testing, sac) occurred between 08:00 and 14:00, except for juvenile social play testing which occurred between 18:00 and 21:00. For Experiment 3, rats were placed on a reversed light cycle to permit cognitive testing during the active phase and food restricted to 85% ad libitum weight. Brains were stored at -20°C in cryoprotectant until staining. TBI intensity was confirmed by righting times and visual inspection of the injury site. Both Sham and Naïve control groups were included to account for the potential impact of surgery on the developing neuroimmune system. All histochemistry utilized well established, vendor-certified antibodies. All equipment and analytical reagents used to perform experimental manipulations and measures are widely available (see Methods for specific vendor information). The statistical tests used were based on the assumptions of normality and homogeneity of variances. The sample sizes and degrees of freedom reflect the number of independent animals. Outliers were determined by values that fall outside the mean ± 1.96 times the standard deviation. For Experiments 1 and 2, the statistical approach was decided a priori to first check for sex differences using Two-Way ANOVAs, followed by One-Way ANOVAs on data collapsed by sex in cases where no sex or interaction effects were present. Correction for multiple comparisons was performed using Tukey tests. For Experiment 3, the complex nature of the RGT made this study too underpowered to detect sex differences. Thus, primary analyses were run on data collapsed by sex, followed by exploratory analyses to probe for potential sex differences, with the caveat that they are merely exploratory at this stage. See Supplementary Table 1 for full statistical details. All data and code presented here are available from the corresponding author upon reasonable request. The authors agree to publish the manuscript using the Mary Ann Liebert Inc. “Open Access” option under appropriate license.

## Acknowledgements

The authors would like to thank the Center for Brain and Spinal Cord Repair at The Ohio State University Wexner Medical Center for the use of surgery core resources and the fluid percussion injury device.

## Author Contributions

MAS and KML conceptualized the study together. MAS, RB, and AEW conducted surgical procedures. MAS, MRB, JEM, and KMM ran the behavioral experiments. MAS, MYM, EY, AZ, and EG ran the histological analyses. MAS, MYM, MRB, CND, JR, and AW analyzed behavioral and molecular data. MAS, CVH, ONK, and KML participated in interpreting the results. MAS generated figures and wrote the manuscript. MAS and KML edited the manuscript. All authors participated in finalizing the manuscript.

## STATEMENTS and DECLARATIONS

## Ethical Considerations

All experiments compiled with the National Institutes of Health Guidelines for the Care and Use of Animals and were approved by the Ohio State University Institutional Animal Care and Use Committee.

## Consent to Participate

Not applicable

## Consent for Publication

Not applicable

## Data Availability

All data presented here are available from the corresponding author upon reasonable request.

