## Supplementary Information and Supplementary Figures for "Pediatric traumatic brain injury elicits acute neuroinflammation and long-term changes in social, cognitive, and decision-making behaviors in male and female rats"

### Table of Contents

**Supplementary Methods**

**Additional and/or expanded methodological detail for select sections - section numbers correspond directly with those in the main body methods*

- **Sections:** 2.1, 2.3, 2.4.1, 2.4.2, 2.4.3, 2.4.4, 2.5.1, 2.5.2, 2.5.3, 2.5.5, 2.6.2, 2.6.3, 2.6.4

**Supplementary Figures**

- **Supplementary Figure 1**: IBA1 and GFAP by Subregion
- **Supplementary Figure 2**: CD68 by Subregion
- **Supplementary Figure 3**: Mast Cells by Subregion
- **Supplementary Figure 4**: Additional Behavior Battery Results
- **Supplementary Figure 5**: Exploratory RGT and FosB Results

**Supplementary Table** (as *separate Excel file*):

- **Supplementary Table 1**: Detailed Statistical Results

### SUPPLEMENTARY METHODS

#### 2.1 Subjects

Timed-pregnant female Sprague Dawley rats were purchased from Inotiv and single-housed in individually-ventilated cages with corncob bedding, with food and water available *ad libitum*. Starting at gestational day 20, cages were checked daily for the presence of pups. Postnatal day (P) 0 was recorded as the day of birth on the first day pups were present in the cage. Pups were sexed between P0-3 but otherwise left undisturbed until P15 when TBI manipulations took place (see below). Pups were then monitored daily as part of postoperative care until either euthanasia (P18) for acute histology assessment cohorts or weaning (P22-24) for behavioral cohorts. The colony room was temperature and humidity-controlled and maintained on a 12-h light-dark cycle (06:00 lights on, 18:00 lights off). For Experiment 2 (see below), rats were transferred to another building containing operant equipment, switched to a reverse light-dark cycle (06:00 lights off, 18:00 lights on), and started on food restriction (maintained ~80% *ad libitum* weight) on P65. All experiments complied with the National Institutes of Health Guidelines for the Care and Use of Animals and approved by the Ohio State University Institutional Animal Care and Use Committee.

#### 2.3 TBI Procedures

On P15, which is roughly equivalent to toddler-age in humans[20,38], male and female pups from each litter were randomly assigned to one of three groups: Naïve, Sham, or TBI (1-3 pups/sex/litter/group). Naïve pups were left undisturbed with the dam, sham pups received all surgical procedures outlined below except for the injury, and TBI pups received all surgical procedures and the injury. The inclusion of both Sham and Naïve control groups allow for the dissociation of effects of injury specifically from any nonspecific effects of early life anesthesia and surgery, which can potentially induce neuroinflammation and/or neurodevelopmental alterations[40].

TBI consisted of a lateral fluid percussion injury (LFPI) (Fig 1A), using a protocol modified from mouse studies and optimized on P15 rats[40]. Briefly, pups were removed from the maternal dam in their home cage, anesthetized with 4% isoflurane, and placed in a stereotaxic frame. The skull was exposed, and a punch biopsy (3mm) was used to trephine a craniectomy in the right parietal bone, midway between bregma and lambda, to reveal the intact dura mater. A hub made from a modified LeurLoc needle was then secured over the craniectomy using cyanoacrylate. Pups were removed from the frame and placed together in a recovery cage on a heating pad for 1-4 hrs. Pups were then anesthetized for 4 mins, and their hubs were attached to the end of the fluid percussion device, which uses a hammer to deliver a fluid pulse (2 atm) over the dura mater exposed during the surgery. TBI pups received this fluid pulse, while Shams were attached to the device but did not get the pulse. The hub was then quickly removed, the skin stapled shut, and righting time (i.e. time to right from a supine position) was recorded. After pups were awake, they were placed back in the home cage with the dam and Naive littermates. Postoperative condition and body weight were monitored on a daily basis for up to a week after surgery, pending the experiment's endpoint.

#### *2.4.1 Tissue Collection*

At 3PDI Naïve, Sham and TBI rats were administered an overdose of sodium pentobarbital and transcardially perfused between 09:00 and 12:00 with 1x phosphate buffered saline (PBS) followed by 4% paraformaldehyde in PBS, pH 7.4. Brains were collected and postfixed overnight in 4% paraformaldehyde at 4°C, then transferred to 30% sucrose in PBS with 1% sodium azide and stored at 4°C. Brains were cut into serial 30μm coronal sections using a sliding microtome, and sections (1/6) were collected into tubes containing cryoprotectant solution and stored at -20°C.

#### *2.4.2 IBA1 and GFAP Staining and Analysis*

One series per animal was removed from cryoprotectant and mounted. Slides were rinsed 3x5 min with PBS then placed in 50% methanol for 30 min to quench background fluorescence. After another 3x5 min in PBS, slides were placed in Tris-EDTA for antigen retrieval, heated to 90°C in a water bath for 10 min, and allowed to sit at room temperature for 10 min. Slides were rinsed 3x5 min in PBS, permeabilized for 1 hr in PBS with 0.4% Triton X100 (PBST), and blocked for 1 hr in PBST with 5% normal donkey serum (NDS). Slides were transferred to primary antibody (Rabbit anti-IBA1 [Wako, 0191741, 1:500]; Mouse anti-GFAP [Sigma, G3893, 1:1000]) in PBST containing 2.5% NDS, and allowed to incubate overnight at 4°C. The next day, slides were rinsed 3x20 min in PBS before secondary antibodies (Donkey anti-rabbit 647+ [Invitrogen, A32795, 1:200]; Donkey anti-mouse 555 [Invitrogen, A32773, 1:333]) were applied in PBST with 2.5% NDS. Slides were allowed to incubate for 2 hrs at room temperature in the dark. Following a final 3x10 min rinse in PBS, slides were incubated with DAPI (Sigma; D9542; 1:10,000) in PBS for 5 min, then coverslipped with Prolong Diamond Antifade Mountant (Invitrogen; P36970).

#### *2.4.3 Microglial CD68 Staining and Analysis*

Upregulated CD68 (phagocytosis marker) expression within microglia can be indicative of a more functional response to an insult such as TBI[28,43,44]. As such, one series was stained with a dual-label IBA1 and CD68 (Mouse anti-CD68, Bio-Rad, MCA341R, 1:500) using the same protocol as 2.4.2.

#### *2.4.4 Mast Cell Staining and Analysis*

Mast cells can be visualized in the brain using a toluidine blue stain, as it binds to their acidified vacuoles[33,45,45]. One series per animal was mounted and stained with 0.5% Toluidine Blue O

(Fisher; BP10710) in 60% acidified ethanol (pH=2.0) for 10 min, followed by sequential washes in ascending alcohols (50%, 70%, 95%, 100%), defatting with xylenes, and coverslipping with Permount (Fisher; SP15100).

A Zeiss Axioimager M2 microscope was used to visualize stained mast cells at 20-40x, and Steroinvestigator was used to tabulate mast cell numbers and degranulation by a blinded observer. Cells were characterized as either granulated or degranulated based on their color, density, and sphericity, with lighter, less dense, and less round cells being degranulated (examples in Fig 4C,D). Mast cells were counted across all sections of the HPC, thalamus, and perivascular space between the two, as these are the main regions where mast cells are located at this age. Counts for total, granulated, and degranulated mast cells are presented as average number per section. HPC and perivascular mast cells were combined for these metrics.

#### *2.5.1 Juvenile Social Play*

Rats were weaned into pairs on P22-24, with sex- and treatment-matched as closely as possible. Juvenile Social Play testing was conducted on P28-P30 [13-15 DPI], as this often represents the peak of play in rats and multiple testing days allow for habituation[40,48]. Play testing was conducted at the start of the dark phase (06:00) under red light after a 4 hr social isolation from their cagemate, in order to capture higher levels of interaction elicited upon reunion with a familiar social stimulus. Home cage pairs of rats were placed back together after this isolation period in a plexiglass box (19x14.5x12 in) and recorded from above and the side for 20 min. Testing order was balanced between pairs across the 3 testing sessions. Play behaviors scored by a blinded observer included chasing, rough and tumble play, and pinning. Data are presented as the total number of play behaviors recorded for each animal across each of the three testing sessions. Following play testing, four pairs were rehoused into groups of 3 to better match sex and treatment group, and rats were maintained in these groups of 2-3/cage throughout behavioral testing.

#### *2.5.2 Spatial Y Maze*

On P61 [46 DPI], rats were tested in the Spatial Y Maze. The Y maze has 3 arms (20x5 in each) that stem from a center point, and arms can be blocked off using an insert. Rats were habituated to red light for 30 min prior to testing. The Spatial Y Maze consisted of 2 phases: familiarization and testing. For familiarization, spatial cues (i.e., white cardboard sheets with distinct black symbols) were placed around the Y maze and one arm was blocked off. Rats were placed in one of the other arms ("starting" arm) and allowed to explore for 10 min. Arms were counterbalanced across groups. After a 1 hr delay, the block is removed and rats are placed back into the starting arm and allowed to explore the newly opened ("novel" arm) and previously opened ("familiar" arm) for 10 min. Videos were recorded from above and time spent in each arm, as well as locomotion, was quantified using Ethovision 11. A discrimination index was calculated using the following formula: (novel duration – familiar duration) / (novel+familiar duration). A more positive index indicates a greater preference for the novel arm.

#### *2.5.3 Spontaneous Y Maze*

Spontaneous Y Maze testing was run on P64 [49 DPI]. Rats were habituated to red light for 30 min prior to testing. The Y maze did not have any arms blocked or spatial cues for this test. Rats were placed in one of the arms (counterbalanced) and allowed to freely explore for 8 min. Videos were scored by a blinded observer for the sequence of entries into each arm. Spontaneously alternating entries are those in which rats progress through the sequence of arms without repeating one (e.g., "1 to 2 to 3" is spontaneously alternating, where "1 to 2 to 1" is not). Data is presented as the percentage of a rat's entries that are spontaneously alternating. Total number of entries is also presented as a gross locomotor measure.

#### *2.5.5 Open Field and Novel Object Recognition*

Rats underwent Open Field and Novel Object Recognition Testing on P70-71 [55-56 DPI]. This two-day test was run under dim white light and consists of 4 phases: habituation (day 1), familiarization 1 (day 1), familiarization 2 (day 2), and novel object testing (day 2). On day 1, rats were placed in the center of an open field (24x24x16 in) and allowed to freely explore for 10 min (habituation). Approximately 2 hrs later, two objects (500 mL bottles filled with blue dyed water) were placed in the open field, and rats were allowed to explore the objects for 10 min (familiarization 1). The next morning, rats were placed back in with the same 2 objects for 10 min (familiarization 2). Four hr later, one of the objects was changed to a 5x4x3 in pink tube rack, and rats were allowed to explore both objects for 10 mins (novel object recognition). Placement of novel and familiar objects were counterbalanced within each group. Ethovision 11 was used to assess center duration, border duration, and distance moved during the first phase (Open Field), as well as time spent exploring (nose within 2 cm) the novel and familiar objects during the last phase (Novel Object Recognition). A discrimination index was calculated using the following formula: (novel duration – familiar duration) / (novel+familiar duration). A positive index indicates preference for the novel object, with higher values indicating higher preference.

#### *2.6.2 Housing and Training*

On P65, rats were transferred to a new housing building for access to operant boxes (Med Associates, St. Albans, VT), switched to an inverted light cycle (06:00 lights off, 18:00 lights on) to enable testing in the active phase, and food restricted to 85% of ad libitum body weight to increase motivation to obtain the sugar pellet reward (45mg F0021, BioServ, Fleming, NJ). Operant testing consisted of daily 30 min sessions (Monday-Friday)[50,51]. From P71-91 [56-76 DPI], rats were habituated to the boxes and gradually taught that nose poking into a lit-up hole yielded a sugar pellet reward. In the latter phase of this training, they underwent a week of forced choice, which familiarized them with the win/loss rules of Choices 1-4 of the RGT (described below). An additional requirement to withhold responding for 5 s prior to choice served as a measure of impulsivity throughout training and the full task.

#### *2.6.3 Rodent Gambling Task Procedure*

Following training, rats entered the free choice RGT, in which they had a choice of those 4 nose poke holes (Choices 1-4) (Fig 6A). Each hole had different probabilities and magnitudes of reward (# sugar pellets) or punishment (timeout with lights on). Choice 2 represented the optimal choice to receive the most pellets. Choice 1 was the safer yet suboptimal choice, yielding consistent but small rewards. Choices 3 and 4 represented gradually more risky decisions, offering more reward but worse punishment. Across several weeks in the RGT, rats should dial into the more optimal Choice 2, choosing it on most trials. Rats were run for 30 sessions (6 weeks). Custom Med-PC IV software was used to record the results of each trail, which were exported to R for analysis. Data were aggregated into 6 5-session bins for analysis and presentation. Analyses were conducted for both the entire testing window (“acquisition”), as well as the final 5 sessions (“stable”).

The primary endpoint of this task is percent choice for each option (Choices 1-4). Overall performance across all 4 choices can also be represented by an RGT Score metric. Score demonstrates the shift from the safer Choices 1 and 2 to the more risky Choices 3 and 4, with higher scores representing more optimal performance overall. The secondary measure of this task is percent of premature responses (i.e. nose poking prior to the hole lighting up), which offers insight into impulsivity and disinhibition. Another endpoint of interest in this cohort is WinStay (i.e. sticking to a previous choice that was rewarded), which offers insight into task optimization. Additional variables RGT yields are Omissions, Number of Trials, Pellets Earned, Choice Latency, Stay, and LoseStay. “Omissions” occur when the rat fails to nose poke while the hole is light. “Number of Trials” is the number of trials a rat manages to get through during a 30 min session. “Pellets Earned” is the number of pellets earned during a 30 min session. “Choice Latency” is the time from a hole lighting up to a rat nose poking. “Stay” refers to a rat sticking with a particular choice over back-to-back trials. “LoseStay” refers to a rat sticking to a previous choice that was not rewarded. In regards to what information can be gleaned from these endpoints, Omissions and Choice Latency are indicative of inattention; Number of Trials and Pellets Earned are broader measures of overall performance; and Stay and LoseStay reveal information about task optimization and perseveration under different win/loss conditions.

#### *2.6.4 Tissue Collection and FosB Assessment*

Following completion of the RGT (4 days after the final testing day), rats were sacrificed and brains were collected and sliced as described in Experiment 1. One series was stained for FosB (Rabbit anti-FosB, Abcam, ab184938, 1:500) analysis in regions previously implicated in the RGT[51], namely the PFC [IL and PL] and nucleus accumbens (NAc) [Shell]. A FosB stain was run and images were acquired as described in 2.4.2. Note that given the extended timing of sac relative to the last testing day, this antibody would exclusively be detecting the delta isoform of FosB. The ImageJ counting tool was used to quantify the number of FosB+ cells by a blinded observer.

### SUPPLEMENTARY FIGURES

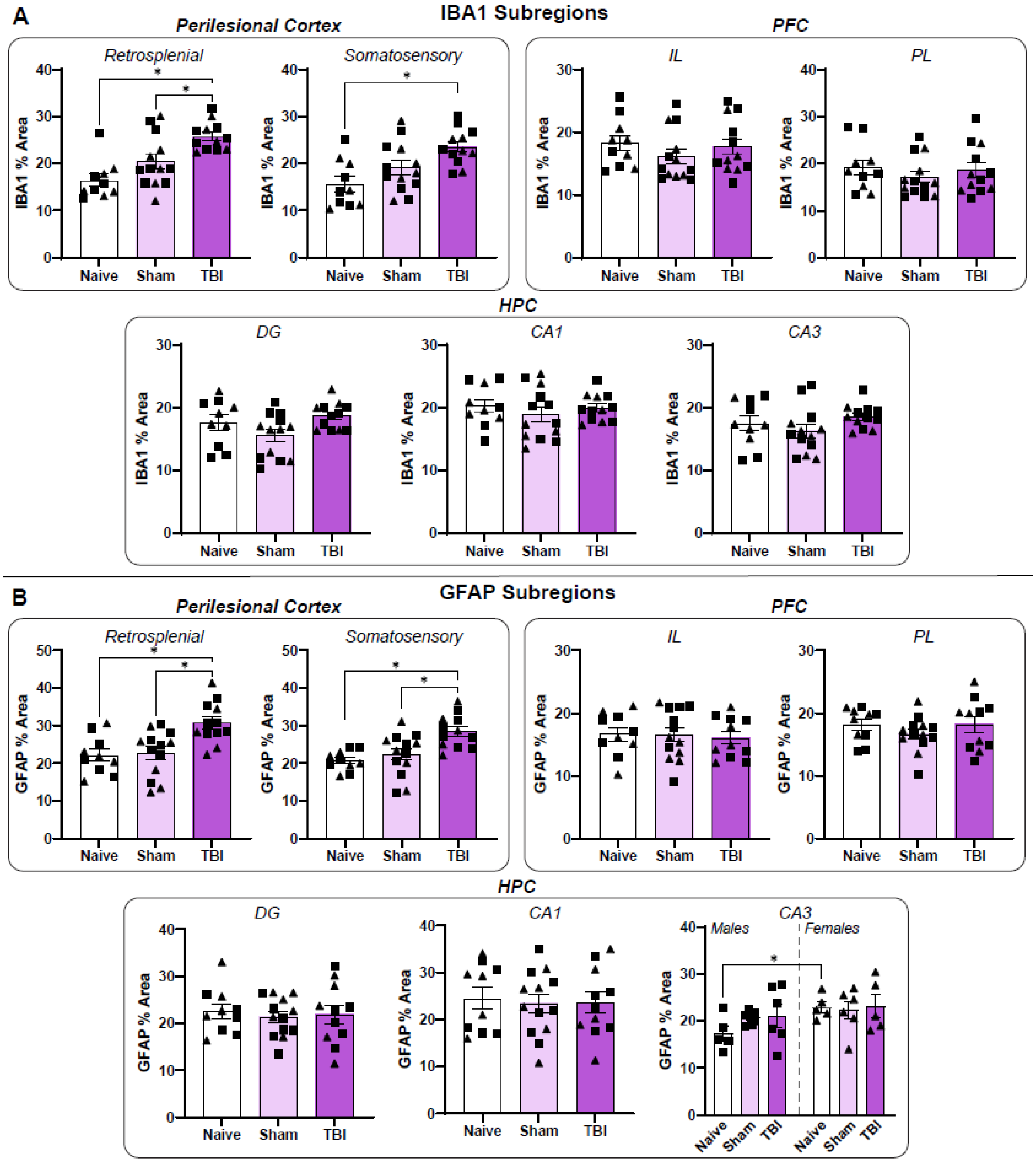


**Supplementary Figure 1: IBA1 and GFAP by Subregion.** (A) TBI increased IBA1 percent area in the retrosplenial and somatosensory Perilesional Cortex, but no subregions of the PFC or HPC. (B) TBI increased GFAP percent area in the retrosplenial and somatosensory Perilesional Cortex, but no subregions of the PFC or HPC. *$p<0.05$. N=5-7/sex/group. Error bars= Mean±SEM. Males are denoted by squares, while females are denoted by triangles.

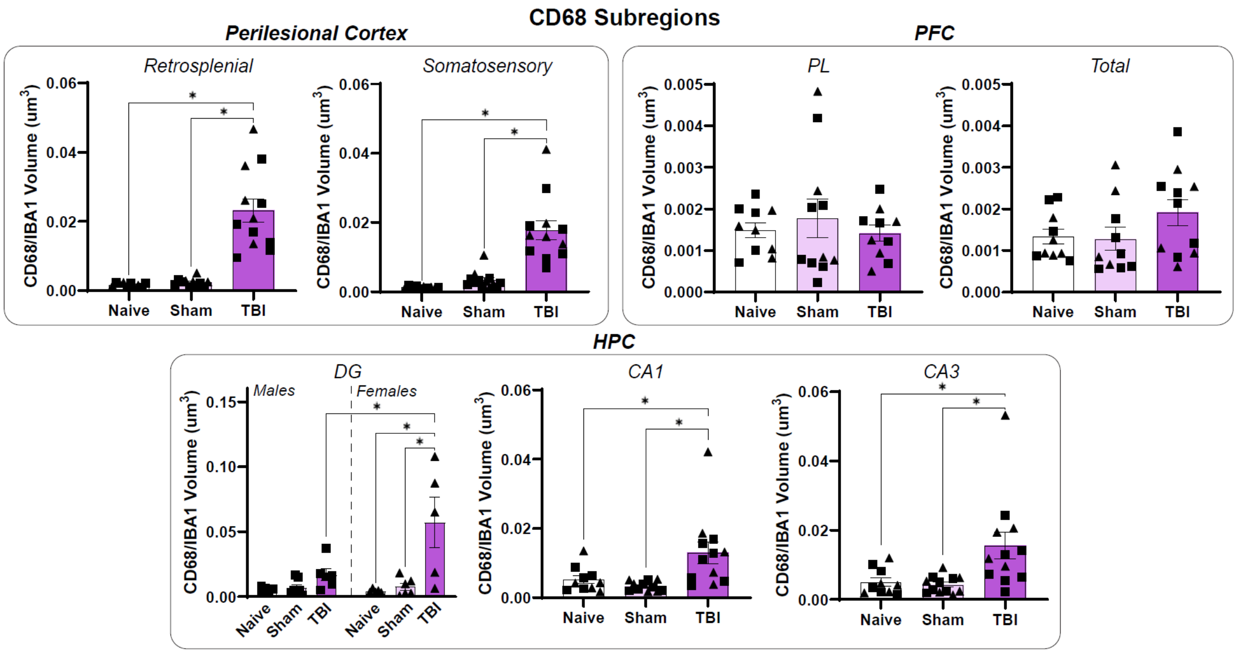


**Supplementary Figure 2: CD68 by Subregion.** TBI increased CD68 expression in microglia in the retrosplenial and somatosensory Perilesional Cortex, as well as the DG, CA1, and CA3 of the HPC. The PL and Total PFC did not exhibit the increase seen in the IL. A sex difference was also present in only the DG HPC, with females exhibiting higher CD68 increases than males. *p<0.05. N=4-7/sex/group. Error bars=Mean±SEM. Males are denoted by squares, while females are denoted by triangles.

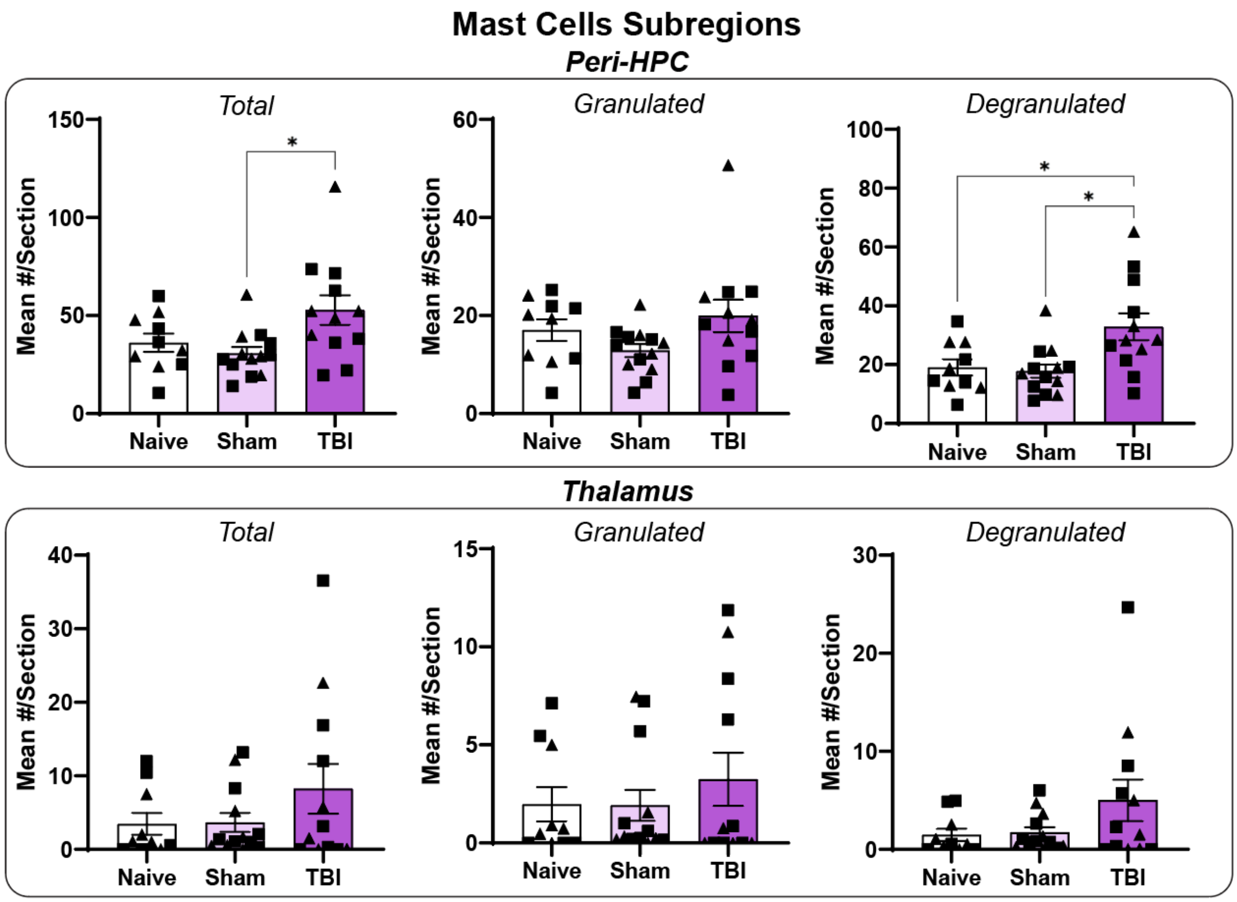


**Supplementary Figure 3: Mast Cells by Subregion.** TBI increased the total number of mast cells in the Peri-HPC region (HPC+velum interpositum). This increase was driven by degranulated, rather than granulated, mast cells. These differences were not significant in the Thalamus. *p<0.05. N=5-7/sex /group. Error bars=Mean±SEM. Males are denoted by squares, while females are denoted by triangles.

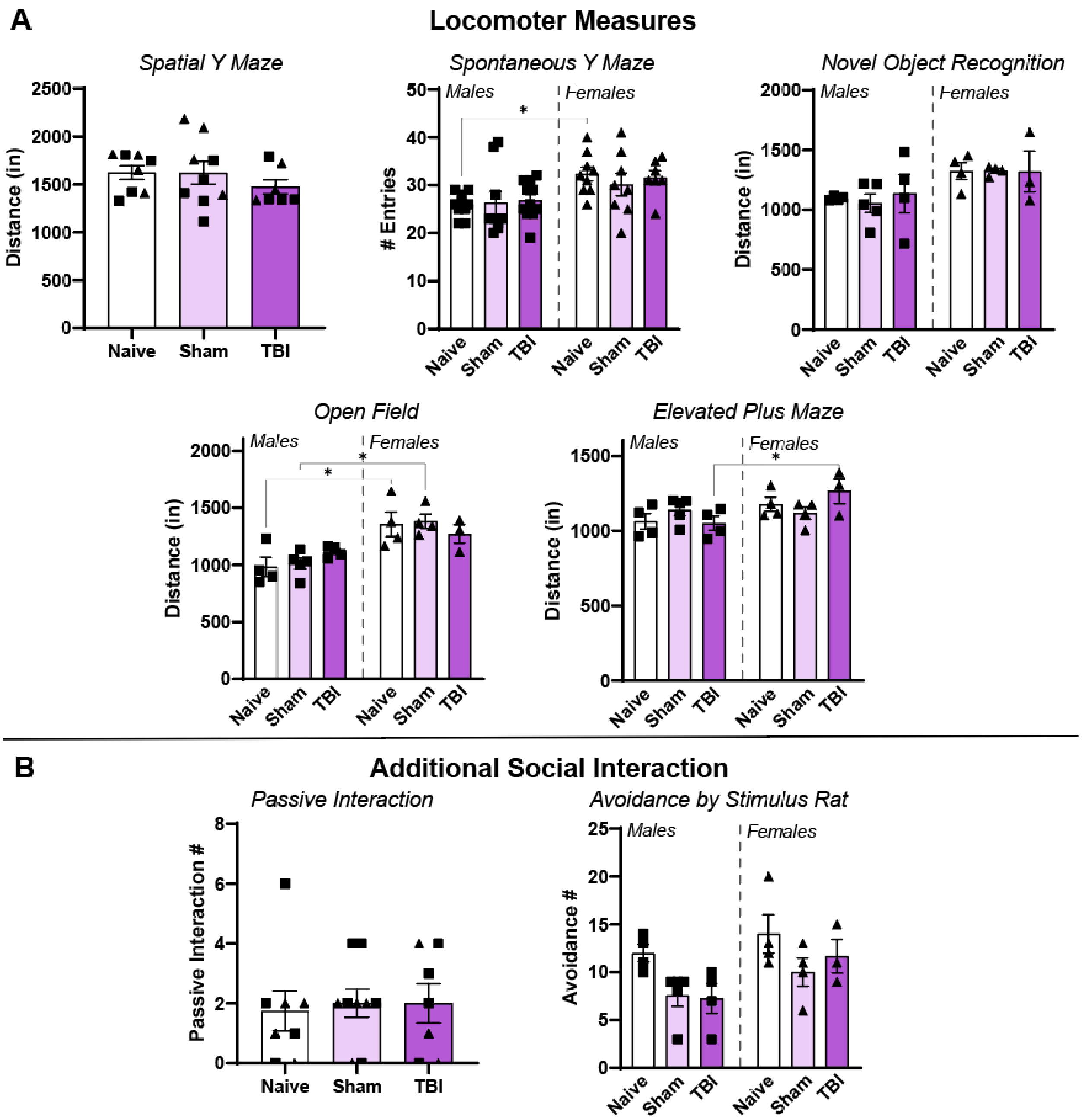


**Supplementary Figure 4: Additional Behavior Battery Results.** (A) Females generally showed higher locomotion across various behavioral tasks than males, although there was some variation by group. (B) TBI did not impact passive social interaction or avoidance by the stimulus rat, although females showed more avoidance in general. *p<0.05. N=3-5/sex/group (ex 8-11 for Spontaneous Y Maze). Error bars=Mean±SEM. Males are denoted by squares, while females are denoted by triangles.

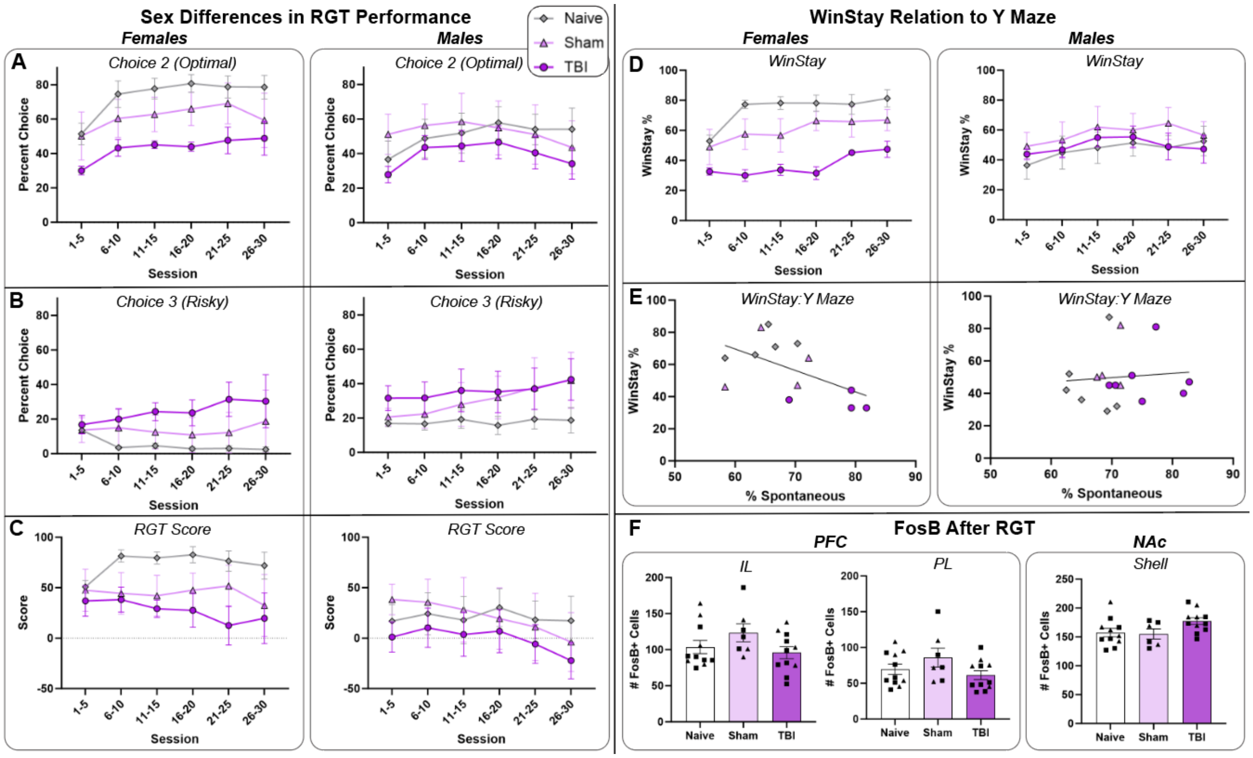


**Supplementary Figure 5: Exploratory RGT and FosB Results.** Including sex as a factor improved Bayesian model fit, suggesting that sex differences were present in RGT performance. (A-B) This difference appears to stem from Naïve females showing a stronger preference for the optimal Choice 2 over the risky Choice 3 than Naïve males, resulting in a higher magnitude shifts in decision-making following TBI in females. (C) Females also exhibited higher Scores in general, as well as a larger delta between Naïve and TBI, supporting the presence of sex differences in RGT performance. (D) The TBI-related WinStay reduction also was driven by females. (E) WinStay in females (but not males) was negatively correlated with Y Maze % Spontaneous, suggesting that the enhanced switching behavior seen in both tasks may be related in females. (F) FosB staining in the PFC and NAc after the RGT did not show any differences in Sham or TBI rats. *$p<0.05$. N=4-7/sex/group. Error bars=Mean±SEM. Males are denoted by squares, while females are denoted by triangles.

**SUPPLEMENTARY TABLE (as separate Excel file)**

**Supplementary Table 1: Detailed Statistical Results.**

*Tab 1: Main Body Figures*. Statistical results for Figures 1-5.

*Tab 2: Supplementary Figures*. Statistical results for Supplementary Figures 1-4.

*Tab 3*: RGT Main Body Figure 6. Statistical results for Figure 6 and additional RGT variables.

*Tab 4*: RGT Supplementary Figure 5. Statistical results for Supplementary Figure 5.
